# KATANIN promotes cell elongation and division to generate proper cell numbers in maize organs

**DOI:** 10.1101/2025.10.05.680529

**Authors:** Stephanie E Martinez, Kin H Lau, Lindy A Allsman, Cassandra Irahola, Cassetty Habib, Isabel Y Diaz, Ian Ceballos, Emmanuel Panteris, Peter Bommert, Amanda Wright, Clifford Weil, Carolyn G Rasmussen

## Abstract

Microtubule severing is essential for proper eukaryotic cell elongation and division. Here we show that the microtubule severing protein, KATANIN, is encoded by two genes in *Zea mays* (maize) called *Discordia3a (Dcd3a)* and *Dcd3b.* Loss of function allele combinations contribute to reduced microtubule severing and decreased cell elongation. The latter results in reduced cell number by slowing entry into mitosis. KATANIN is important for preprophase band (PPB) formation and positioning, and nuclear positioning in symmetric cell divisions. A combination of these defects contributes to generating mutant plants with small size and aberrant shape.

## Introduction

Dynamic microtubule reorganization is critical for development in eukaryotes. Microtubule dynamics are regulated by hundreds of proteins, many of them conserved across the eukaryotic lineage. One critical type of microtubule reorganization occurs via microtubule severing, which is performed by enzyme complexes, including KATANIN, that hydrolyze ATP to sever microtubules. One subunit of the KATANIN microtubule severing complex is an ATPase called p60. The p60 subunit alone has a relatively low binding affinity for microtubules but still forms a multimeric ring capable of ATP-dependent severing^1–3^. p60 is effectively recruited to microtubules by the p80 subunit both *in vitro*^4^ and *in vivo*^5^. In plants, the KATANIN complex assembles primarily at microtubule crossovers^6–8^. The p60 and p80 subunits together form heterodimers that assemble into a higher-order ring complex that grips the *β*-tubulin polyglutamate tail to generate the force for ATP-hydrolysis dependent severing^9^. Katanin interaction with the *β*-tubulin subunit has also been confirmed *in vivo* in *Caenorhabditis elegans*^10^.

In plants, *p60* mutants have been isolated from many forward mutant screens due to their striking vegetative phenotypes^11–15^. In *Arabidopsis thaliana*, *Cucumis sativa* (cucumber), and *Oryza sativa* (rice), the p60 subunit is encoded by one gene^13,15,16^. In *A. thaliana*, the *p60* mutant has aberrant cell elongation, small organs, less cellulose, altered lignin accumulation and disrupted cortical microtubule organization^17–22^. Similarly, the cucumber *p60* mutant has smaller fruits with fewer cells^15^. In rice, the *p60* mutant has small and round leaves, and shorter cells with aberrant cortical microtubule organization^13^. KATANIN also mediates rapid microtubule reorganization in response to environmental cues, affecting morphological changes in response to stimuli such as blue light and mechanical stimulation^23–25^. Despite the striking macroscopic defects, an understanding of how KATANIN contributes to growth has not been fully explored.

In addition to facilitating proper interphase microtubule organization, KATANIN-mediated severing contributes to remodeling microtubule arrays during cell division. The preprophase band (PPB), a land-plant-specific microtubule array, accumulates prior to mitosis at the future division site ^26,27^. The *p60* mutants often generate abnormal PPBs ^28,20,29^. Abnormal microtubule arrangements include frayed PPB formation, where one side of the PPB has less or poorly organized microtubules, as well as delayed PPB narrowing^20,28^. Although metaphase spindles are occasionally tilted ^28,30^, time-lapse imaging showed that misoriented spindles did not lead to altered final division plane positioning in petioles^20^. There are no abnormalities in the early formation of the cytokinetic structure called the phragmoplast, although it too is occasionally misoriented. Later, during phragmoplast expansion, *katanin* mutants form a double-arrow shaped phragmoplast, also observed in mutants lacking other microtubule-binding proteins^31,32^, likely due to aberrant unseverable microtubule connections between the reforming nuclear envelope and phragmoplast microtubules^20,28^. Even though PPB narrowing, mitosis, and cytokinesis are slower in *katanin* mutants, no obvious division orientation defects are observed in epidermal petiole cells^20^. In contrast, ectopic or misoriented divisions in roots lead to altered cell fates affecting protoxylem, pericycle and root hair cells^14,28,30,33^. Whether misoriented divisions in *katanin* mutants are due solely to misoriented PPBs or additional phragmoplast guidance defects is currently unclear^20,28,30,34^.

Recent focus has further highlighted connections between the nucleus and the PPB, and consequently the final location of the division^35,36^. The MAIZE LINC KASH AtSINE-LIKE2 (MLKS2) protein tethers the nucleus to the actin cytoskeleton. MLKS2 plays a critical role in chromosome segregation during meiosis, maintaining proper nuclear morphology, and asymmetric division plane positioning ^37^. Further, it contributes both to nuclear polarization via movement towards the correct location and the maintenance of nuclear positioning prior to subsidiary cell divisions in maize ^38^. The nucleus polarizes before the PPB forms^39,40^, often dependent on an intact actin^41^ or microtubule cytoskeleton^42^. When the nucleus is polarized incorrectly or is not maintained near the future division site, the PPB forms in the incorrect location, leading to misoriented divisions^37,38^. Formation of incorrectly positioned PPBs and unpolarized nuclei occur in subsidiary cell divisions in mutants lacking actin nucleators, suggesting that connections between actin and the nucleus are important for division positioning ^43,44^. Similarly, when the nucleus is displaced by centrifugation after formation of a PPB, an additional new PPB forms near the nucleus^45^. Together, these data highlight the important role of the nucleus in positioning the PPB, but we show here that the PPB also positions the nucleus.

Here we characterize the function of two *Katanin p60* homologs in maize, and define their contributions to plant and organ growth. We show that cell elongation defects and reduction in cell numbers together account for smaller organs in the *p60* double mutant, assessed via analysis of growing roots and leaves, as well as fully grown leaves. Further, the *p60* mutants have defects in PPB formation which are correlated with defects in nuclear positioning, suggesting that KATANIN-mediated severing alters nuclear positioning. Finally, symmetric division plane positioning defects are due to occasionally misoriented PPBs, not phragmoplast guidance defects.

## Results

### Identification of *katanin* p60 mutants in *Zea mays*

The maize genome contains two *Katanin p60* genes, Zm00001eb156560 (*Dcd3a* on Chromosome 3) and Zm00001eb360490 (*Dcd3b/clt1* on Chromosome 8): individually they are mostly redundant but loss of both alters proper growth and division. The *discordia3* (*dcd3*) mutant was identified by subsidiary cell division positioning defects in the leaf epidermis, similar to *dcd1* ^46,47^ and *opaque1/dcd2* ^48–51^. While originally thought to be a single mutant, genetic analysis revealed that *dcd3* was a double mutant in both *Dcd3a* and *Dcd3b.* DNA sequencing identified a 1 base pair deletion in Zm00001eb360490 on Chromosome 8, *dcd3b*, which led to a frameshift induced premature stop codon towards the C-terminus of the protein (Figure 1A). Additionally, *dcd3* also contained several missense mutations mapped to Chromosome 3 at the Zm00001eb156560 locus, *dcd3a-1* (Figure 1A). The deleterious *dcd3a-1* allele is identical to the allele from the maize inbred line CML228. From an independent EMS screen, a semi-dominant mutant, *Clumped tassel1* (*Clt1*), was identified by its short stature and clumped tassel phenotype and was mapped to an interval on the long arm of Chromosome 8^52–54^. Subsequent characterization identified a missense mutation (C to T at coding sequence position 1061), changing amino acid 354 from serine to phenylalanine within the ATPase domain (Figure 1A, Supplementary Figure 1). *Clt1* homozygotes are shorter than wild-type siblings and contain cells that are more isotropic and often have division plane orientation defects (Figure 1E). *Clt1/+* plants have an intermediate plant height (Figure 1C).

**Figure 1.**
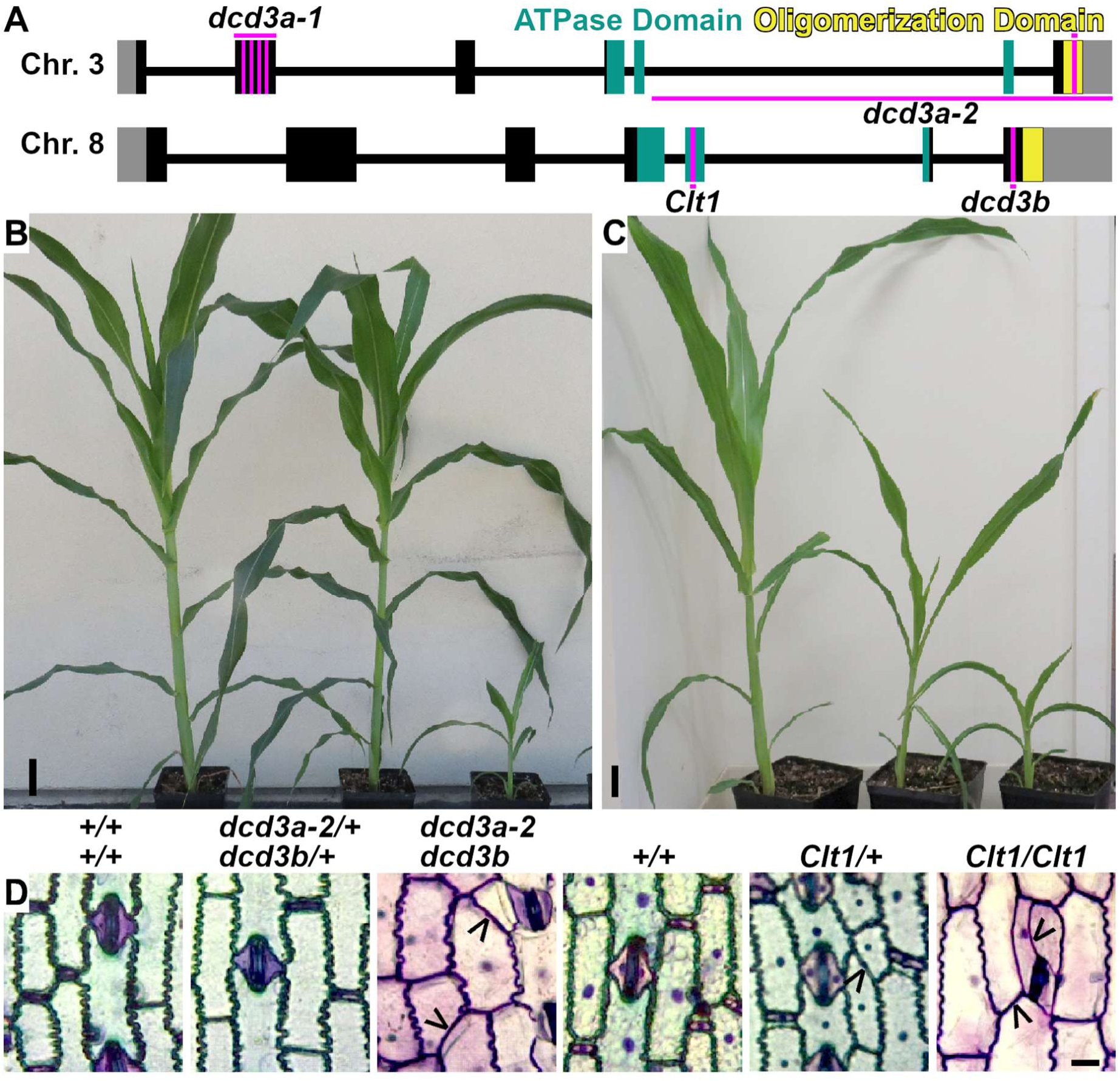
Two *KATANIN p60* encoding genes in maize with two independent mutant alleles each. (A). Gene schematic depicting the position of SNPs compared to B73. (B, C). Representative 4- or 5-week-old sibling plants. Scale bar = 5 cm. (D, E). Toluidine-blue O stained epidermal peels of the eighth leaf. Black arrowheads indicate likely division plane orientation defects. Scale bar = 10 µm.

The *Clt1/+* mutant was crossed to the Nested Association Mapping (NAM) parent lines to identify *Clt1/+* enhancers. Two enhancers that significantly reduced *Clt1/+* plant height were from maize inbred lines CML228 and CML333 (Supplementary Figure 2A and 2B). The genic region containing the enhancers mapped to Chromosome 3, overlapping the location of *Dcd3a*. As described above, CML228 contains an alternative allele of *Dcd3a (dcd3a-1).* CML333 contains a deletion of ∼ 3Kb in Zm00001eb156560, this mutant allele is, henceforth, termed *dcd3a-2* (Figure 1A, Supplementary Figure 2E and F). Homozygous *dcd3a-2 Clt1* double mutants did not germinate, whereas *dcd3a-2 dcd3b* double mutants were viable. However, *dcd3a-2 dcd3b* double mutants were sterile and had short stature and division positioning defects. Mutants also had reproducible differences in TBO staining (Figure 1D and E), suggesting altered cell-wall composition, which may be similar to cell-wall alterations in Arabidopsis *katanin* mutants^17^. Due to the complexity of possible mutant allele combinations, we focused on the likely loss-of-function *dcd3a-2 dcd3b* double mutants.

### *p60 katanin* mutants have decreased microtubule severing frequency

To investigate the contributions of p60 KATANIN on microtubule severing, four-week-old plants expressing a live-cell marker for microtubules were used to image microtubules in the leaf elongation zone. First, we analyzed cortical interphase microtubule density and anisotropy (directional uniformity) in mutant and wild-type cells: microtubule density was similar in all genotypes (Figure 2A and B), while anisotropy was significantly lower in *dcd3a-2 dcd3b* cells (Figure 2C), similar to Arabidopsis *katanin* mutants^6,17,20,55,56^. For microtubule severing, we selected a subset of cells with similar cortical microtubule density and anisotropy to analyze (≥15 cells per genotype, 3 or more plants, Supplementary Figure 3). Microtubule severing frequency was obtained by counting the number of microtubule severing events at microtubule crossovers per area over time (Figure 2D). Wild type and double heterozygote *dcd3a-2/+ dcd3b/+* plants had similar severing frequencies (Figure 2E), similar microtubule density and anisotropy and grew similarly. This, together with sterility of the *dcd3a-2 dcd3b* mutant plants, prompted us to use the double heterozygote as a wild-type control in subsequent experiments as the *dcd3a-2 dcd3b* double mutants were most often generated from *dcd3a-2/+ dcd3b X dcd3a-2 dcd3b/+* or reciprocal crosses.

**Figure 2.**
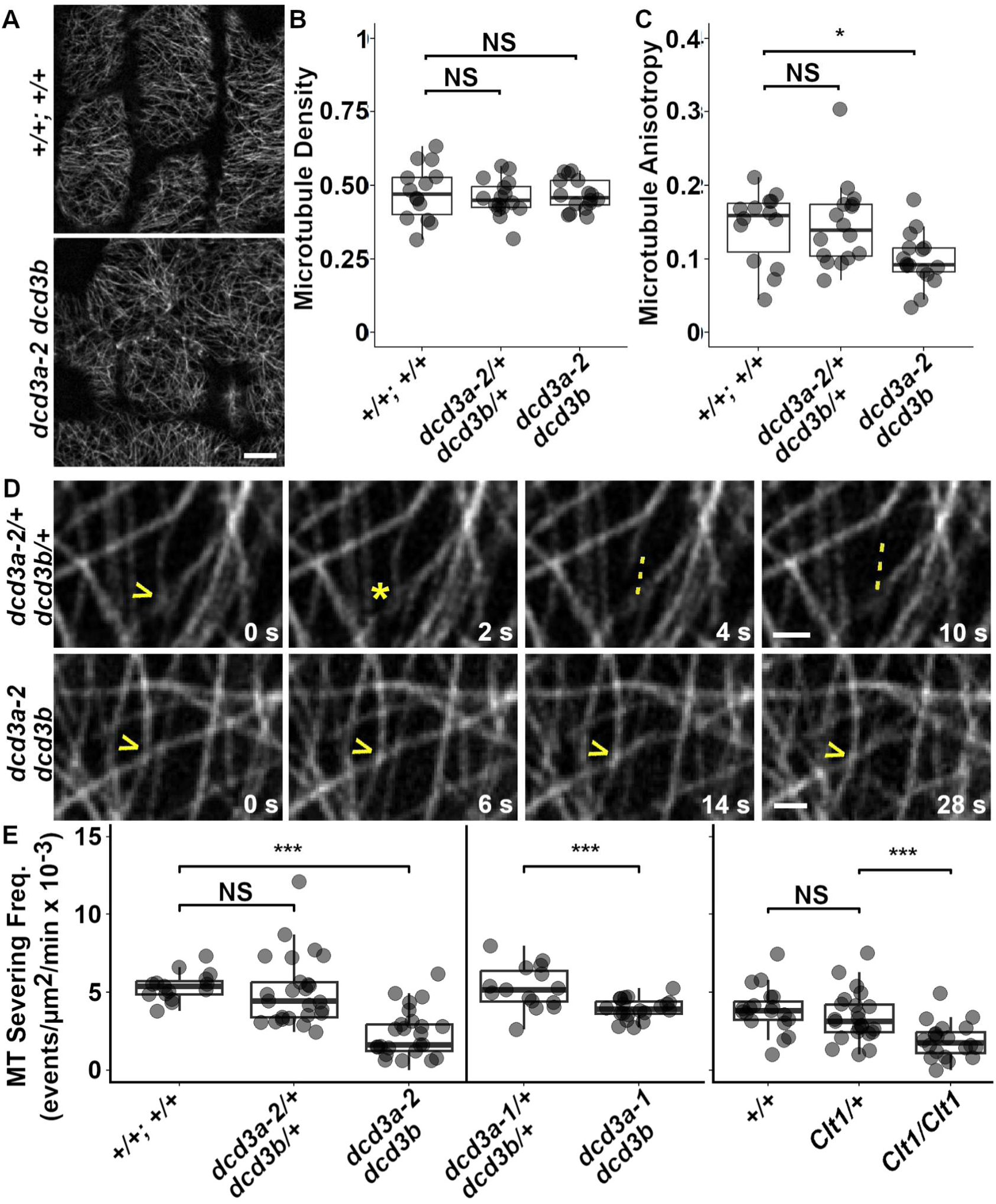
*Katanin p60* mutants have reduced microtubule severing compared to wild-type siblings. (A). Representative micrographs of interphase cortical microtubules. Scale bar = 10 μm. For microtubule density (B) and microtubule anisotropy (C), 14 or more cells from 3 plants per genotype were analyzed. (D). Representative micrographs showing microtubule severing in plants expressing YFP-TUBULIN. Yellow arrows = microtubule crossover. Yellow asterisk = severing event. Yellow dotted line follows the severing-induced depolymerization. Scale bars = 1 µm. (E). Boxplots of microtubule severing frequencies, each dot represents one cell. 15 or more cells from 3 or more plants per genotype were analyzed. Student’s t-test, p-value NS > .05, * = 0.01, *** < 0.001.

Both *dcd3a-2 dcd3b* double mutants and homozygous *Clt1* mutants had reduced microtubule severing compared to wild-type siblings. The *dcd3a-2 dcd3b* double mutants had >2-fold reduction in microtubule severing frequency when compared to wild-type siblings (Figure 2B; T-test p-value < 0.001). Additionally, *dcd3a-1 dcd3b* double mutants had a >1.3-fold reduction in microtubule severing frequency when compared to wild-type siblings (Figure 2B, T-test p-value = 0.0026). Surprisingly, there was no significant difference in microtubule severing frequency between *Clt1/+* and wild type elongation-zone cells (Figure 2B; T-test p-value = 0.33). This was unexpected because the *Clt1/+* phenotype is macroscopically obvious, although less severe than double mutants or *Clt1* homozygotes. *Clt1* homozygous mutants had >2-fold reduction in microtubule severing frequency when compared to wild type, similar to the *dcd3a-2 dcd3b* double mutant (Figure 2B; T-test p-value < 0.001). While severing frequencies of several other mutant combinations tested were indistinguishable from wild-type, single *dcd3b* homozygous mutants were ∼1.3-fold lower than wild-type siblings (Supplementary Figure 4). Together, these data indicate that both p60 proteins significantly contribute to microtubule severing but that DCD3B plays a more prominent role, particularly in the elongation zone of adult leaves where severing was measured.

### The *dcd3a-2 dcd3b* double mutants have small roots and leaves

Similar to *katanin* mutants in other plants^17,57–59^, *dcd3a-2 dcd3b* double mutants have reduced overall height (Figure 1B) and ∼3-fold smaller roots at day 4 with slower growth compared to wild-type siblings (Supplementary Figure 5). We assessed V5 leaf elongation rates (n ≥ 15 plants each) and while wild-type leaf elongation rates were similar to previous reports^60–62^ (2.8 mm/hr ± 0.2 mm SE), mutants elongated significantly slower (1.6 mm/hr ± 0.01 SE). Additionally, mutants also had smaller fully-grown leaves (Figure 3). First leaves of 11-day-old mutant plants were ∼3.5-fold smaller than corresponding wild-type sibling leaves (V1, Figure 3C, Supplementary Table 1). Further, the *dcd3a-2 dcd3b* double mutants showed ∼3-fold smaller leaves compared to wild-type leaves at other developmental stages: fully-expanded 5th leaves (V5, transitioning between juvenile and adult leaves) and fully-expanded adult leaves (V8) (Figure 3F, I, Supplementary Table 1). The ∼3-fold smaller leaves and roots, and slower growth rates thus account for the smaller size of *dcd3a-2 dcd3b* mutants as compared to wild-type siblings.

**Figure 3.**
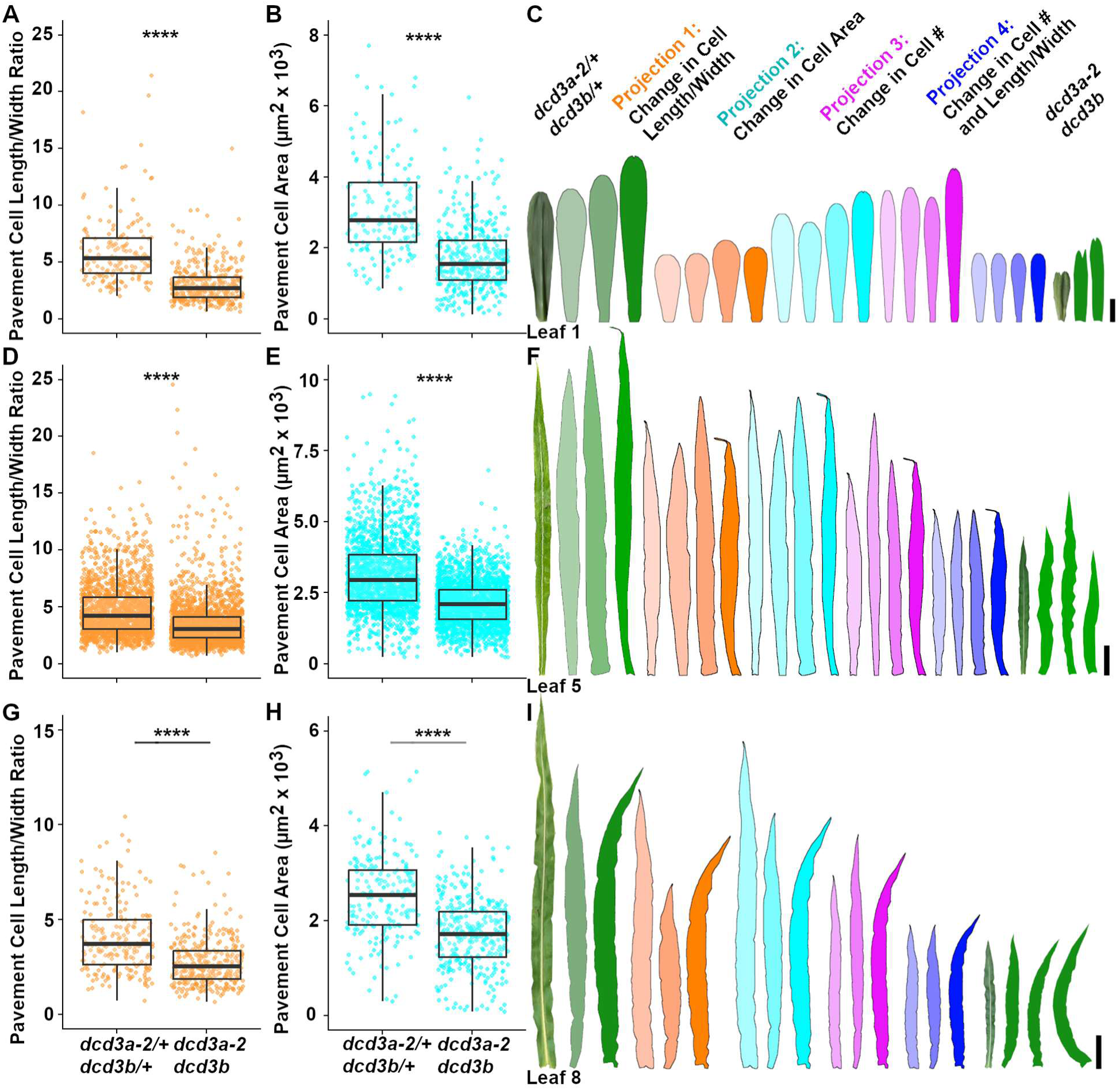
*dcd3a-2 dcd3b* double mutants have smaller leaves due to fewer and smaller cells. (A, D, G) Pavement cell length/width ratio in wild-type *dcd3a-2/+ dcd3b/+* and mutant *dcd3a-2 dcd3b* for (A) leaf 1 (D) leaf 5 (G) leaf 8 (B, E, H) Total cell area (µm^2^ x 10^3^) of pavement cells for (B) leaf 1 (E) leaf 5 (H) leaf 8. N ≥ 3 plants per genotype. Median and quartiles are shown in the boxplot. Whiskers are 1.5X the interquartile range. (C, F, I) Leaf projections for (C) leaf 1, (F) leaf 5, and (I) leaf 8. Scale bars = 1, 5, and 10 cm, respectively. Welch’s two sample t-test p-value < 0.00001 ****. Supporting data in Supplementary Tables 1, 2 and Supplementary Figure 6.

### Small aberrantly expanded cells do not fully account for the small leaves of mutants

To determine why mutants had smaller leaves, we first tested the hypothesis that cell elongation defects caused by decreased microtubule anisotropy generate small leaves. Epidermal pavement cell and stomatal cell complex outlines were obtained from the base, middle, and tip of the leaf to measure cell areas (Supplementary Figure 6, Supplementary Table 2). In leaves one (V1), five (V5), and eight (V8), pavement cells of the *dcd3a-2 dcd3b* double mutant were significantly less elongated (Figure 3A, D, G) and smaller than wild-type siblings (Figure 3B, E, H), similar to previous reports in other plants^17,22,59^. Using cell length and width parameters of the mutants (Figure 3A, D, G), we generated a projection to reflect how this would alter wild-type leaf size (Figure 3C, F, I: Projection 1, orange). Not surprisingly, these projections generated shorter and wider leaves, with a projected leaf area ∼46-77% of wild-type leaves. However, the projections were still ∼1.6-2.4-fold larger than the mutant leaves. This indicates that altered cell elongation partially contributes to the small leaves observed in *dcd3a-2 dcd3b* mutants. Next, we used the average smaller cell areas of the *dcd3a-2 dcd3b* double mutants to generate Projection 2 (Figure 3C, F, I, cyan). Projection 2 generated ∼0.7-0.8-fold smaller projected leaves when compared to wild-type but ∼2.2-2.7-fold larger than the mutants). Projection 1 and Projection 2 explicitly assume that wild-type and *dcd3a-2 dcd3b* mutants have the same number of cells within the leaf.

### *dcd3a-2 dcd3b* mutants have fewer cells than wild-type plants in leaves

Because smaller cell areas did not generate leaf projections that were similar to *dcd3a-2 dcd3b* mutant leaves, we next estimated cell numbers in wild-type and mutant leaves. We estimated the cell number based on extrapolations from the number of cells per area in several locations (see Materials and Methods). *dcd3a-2 dcd3b* mutants had approximately 3-fold fewer cells in leaf 1 compared to wild-type siblings (Supplementary Table 1; T-test p-value = 0.002). Other mutant leaves (V5 and V8) also had ∼2-2.7-fold fewer cells than wild-type siblings (Supplementary Table 1). We next used the average of these reduced mutant cell numbers to generate new projections of wild-type leaves (Figure 3C, F, I: Projection 3, magenta). The projected leaves had ∼42-58% the wild-type leaf area but were still between 143% and 197% larger than the mutant leaves (Figure 3C, F, I). This suggests that reduced cell number contributes to the different leaf sizes. Finally, we generated a projection that combined both cell shape differences and reduced cell number, Projection 4 (Figure 3C, F, I, blue). Not surprisingly, these projections generated leaves more similar to *dcd3a-2 dcd3b* leaves, with projected leaf areas ∼101-114% the size of the average *dcd3a-2 dcd3b* leaves. Taken together, these results suggest that p60 plays a combined role in cell elongation and cell proliferation. Based on the relative reductions in leaf area, we estimated that cell elongation defects contributed slightly more than reduced cell proliferation to overall size.

### *dcd3a-2 dcd3b* mutants have delayed entry into mitosis, but no delay in cell division progression

To investigate whether fewer cells observed in *dcd3a-2 dcd3b* mutants were due to delays in cell division, we performed time-lapse analysis of symmetric cell divisions in four-week-old plants. Contrary to delays observed in Arabidopsis *katanin p60* mutants^20^, transverse symmetric division duration measured from the start of metaphase (spindle formation) through phragmoplast disassembly was similar between *dcd3a-2 dcd3b* double mutants and wild-type siblings, as were metaphase and telophase durations (Supplementary Table 3) Phragmoplast expansion rates were also similar (*dcd3a-2/+ dcd3b/+*: 0.31 µm/min ± 0.02 SE, n = 69 cells; *dcd3a-2 dcd3b*: 0.30 µm/min ± 0.02 SE n = 48 cells; Wilcoxon rank sum test p-value = 0.87). Similar cell division times between the double mutant and wild-type sibling suggest that the reduced cell number is not due to significant delays in metaphase through the end of cytokinesis.

As an independent test, we compared the relative proportions of specific division stages in symmetrically dividing zones (ligule height ∼2 mm) using developmentally matched micrographs of *dcd3a-2 dcd3b* double mutant and wild type plants, assessed with a live cell marker for microtubules in late G2 and actively dividing cells (e.g., late G2/prophase, metaphase, or telophase, Figure 4A). The cell-cycle stages represented in mutants and wild types were proportional to each other (Supplementary Table 4), indicating that *dcd3a-2 dcd3b* double mutants do not have delays in any mitotic stage compared with wild-type siblings. That no particular mitotic stage is overrepresented is consistent with time-lapse imaging showing no significant difference in mitotic progression timing.

**Figure 4:**
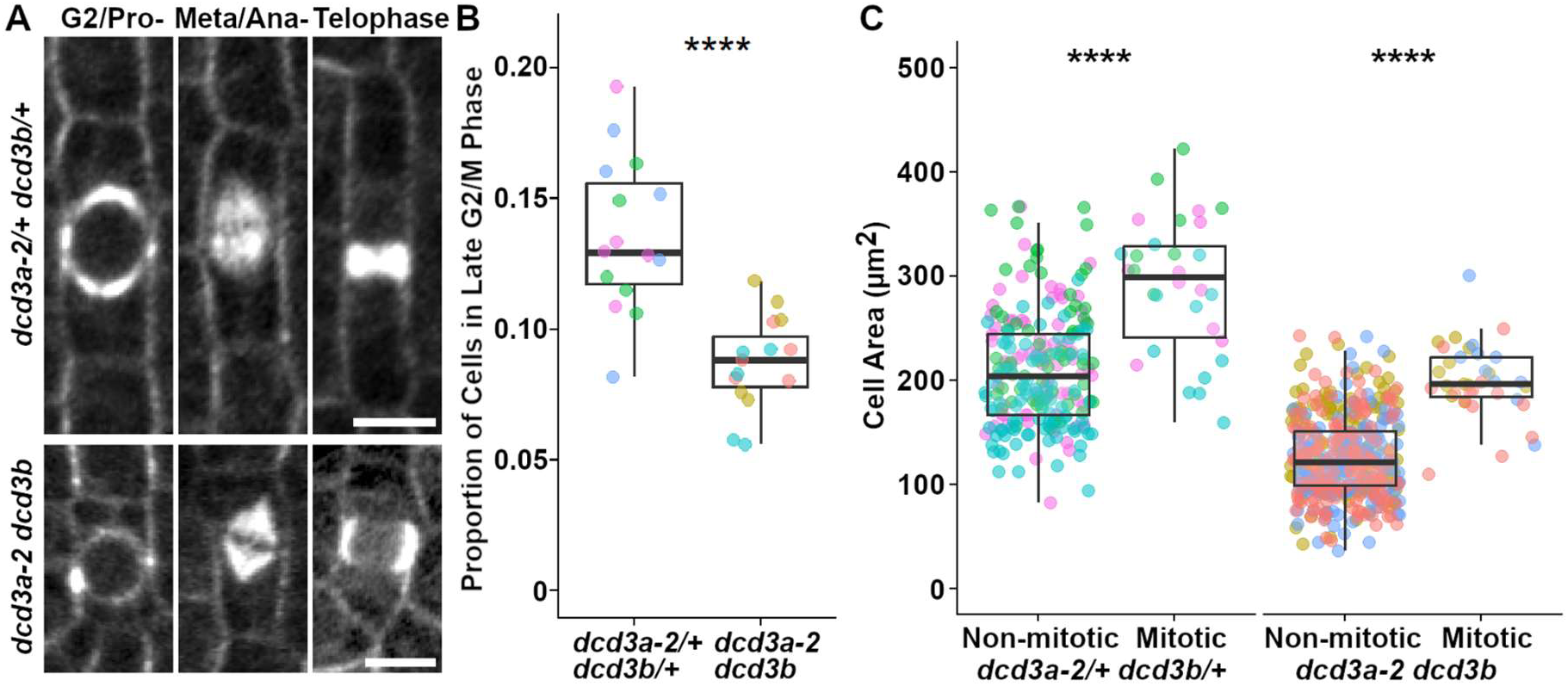
The *dcd3a-2 dcd3b* double mutants have fewer actively dividing cells in the symmetric division zone. (A). Micrographs of cells counted during different cell stages for 3 plants per genotype. *dcd3a-2/+ dcd3b/+* and *dcd3a-2 dcd3b*. Scale bars = 10 µm. (B). Boxplot of the proportion of late G2/M cells, dot represents one micrograph.. (C). Cell area (µm^2^) in wild-type (non-mitotic: 210 µm^2^ ± 4 SE from 226 cells; mitotic: 291 µm^2^ ± 12 SE from 30 cells) and mutant (non-mitotic: 126 µm2 ± 2 SE from 430 cells; mitotic: 199 µm^2^ ± 6 SE from 36 cells). P-values are from Welch’s two sample t-tests. Each color represents an individual plant, N = 3 plants per genotype. Welch’s two sample t-test p-value < 0.00001 = ****. Supporting data in Supplementary Table 4 and 5.

Although the proportions of specific mitotic stages were similar, significantly fewer cells in the symmetric division zone were actively dividing in mutants compared to wild-type plants. *dcd3a-2 dcd3b* mutants have ∼1.6-fold fewer total cells in preprophase/prophase through cytokinesis when compared to wild-type siblings (Figure 4B; Supplementary Table 5). Live-cell imaging and simulation experiments indicate that cell size and cell-cycle progression are linked ^63,64^. To determine whether small cell size might be responsible for delayed entry into mitosis, we measured areas of mitotic and non-mitotic cells from mutant and wild-type dividing cells described above (3 plants each, from V8 and V12 respectively). *dcd3a-2 dcd3b* mutants had smaller cell areas in both mitotic and non-mitotic cells (Figure 4C), with 1.6-fold larger mitotic cells. Wild-type mitotic cells were 1.4-fold larger than non-mitotic cells. As expected, cells entering division tended to be larger than non-mitotic cells. However, since the *dcd3a dcd3b* double mutant cells start small, it is likely that they also enter mitosis after significant delays in G1 or G2, or both, to overcome minimum cell-size thresholds.

### Misoriented PPBs generate misoriented divisions, but abnormally frayed or incomplete PPBs do not alter division plane orientation in symmetric divisions

Similar to *katanin* mutants in other plants^20,28,33^, the *dcd3a-2 dcd3b* double mutant has a high frequency of abnormal PPBs (∼46%, n = 83/182 cells from 3 plants, Table 1). In contrast, most PPBs in wild-type siblings were normal (∼91%, n = 147/161). PPBs classified as “normal” completely encircled the nucleus with consistent CFP-TUBULIN fluorescence intensity across the whole band. Uneven PPBs, where the microtubule accumulation is not consistent across the whole band, occurred in ∼32% of mutants but only ∼5% in wild-type cells. While rarely observed, four-sided, three-sided, and split PPBs occurred with similar frequency in wild-type and mutant cells. The *dcd3a-2 dcd3b* mutant also had ∼9% one-sided or incomplete PPBs, similar to Arabidopsis *katanin* mutants ^20,28^, and 5% misoriented PPBs, where the PPB was positioned perpendicular rather than parallel across in the cell (Figure 5A) to generate a curved division. *dcd3a-2 dcd3b* mutants had ∼46% normal PPBs (Figure 5A). Neither one-sided nor misoriented PPBs were observed in wild-type siblings.

**Figure 5.**
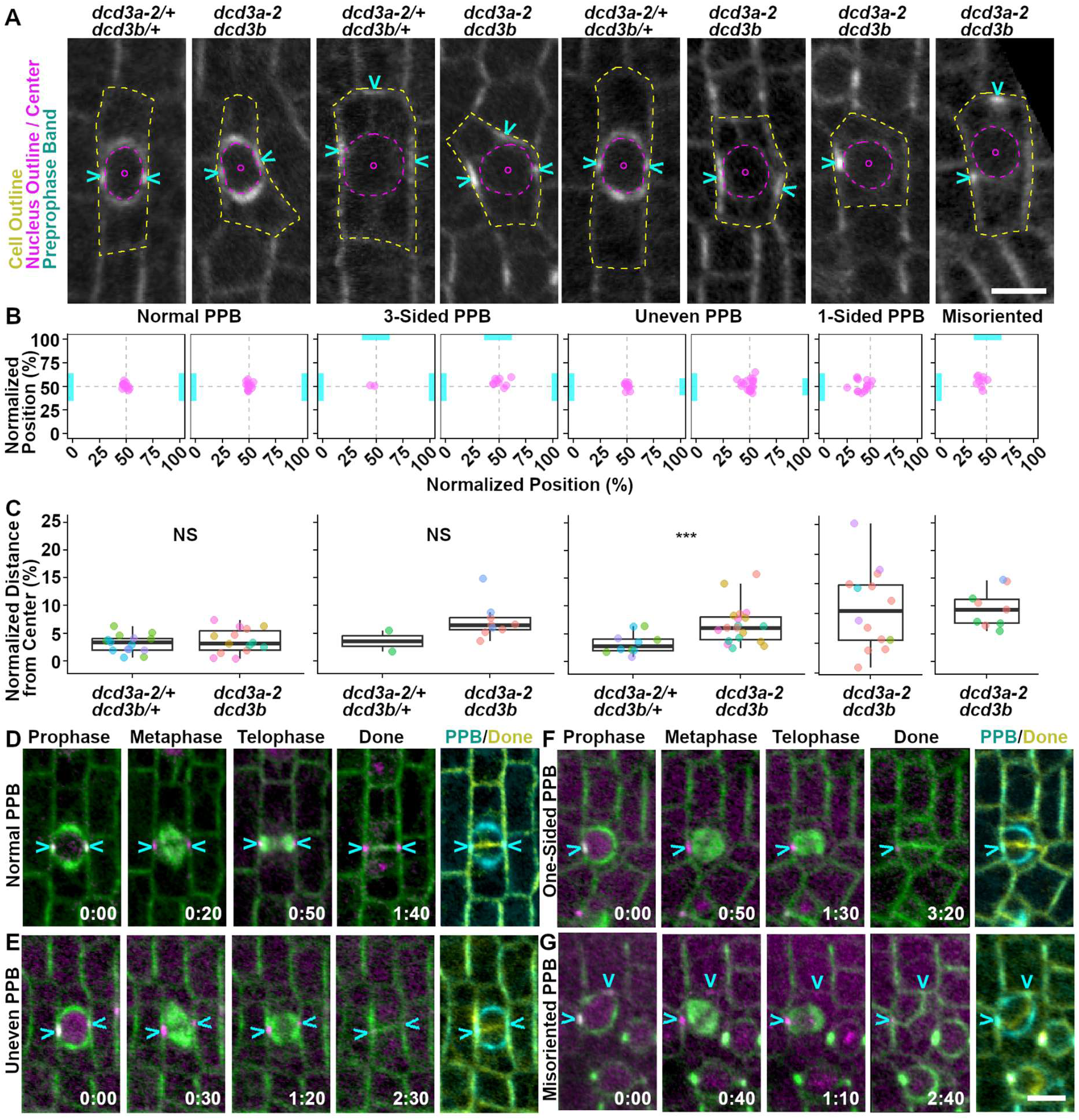
Nuclear positioning defects in *dcd3a-2 dcd3b* mutants may be caused by aberrant PPB accumulation, while misoriented PPBs lead to misoriented divisions. A). Representative micrographs comparing nuclear positioning and PPBs in wild-type and mutant cells. Cell outline = yellow dotted line. Nucleus outline and centroid = magenta. Cyan arrows = PPB. For B and C, 3 or more plants per genotype were analyzed unless mentioned. B). Normalized nuclear position (%). Each dot = one cell. Cyan bars indicate the PPB type and position. C). Normalized distance of the nuclear center to the cell center (%). Each color = a plant. For normal PPBs: n ≥ 14 cells per genotype; Welch’s two sample test p-value = 0.52. Two plants per genotype had 3-sided PPBs; Welch’s two-sample t-test p-value = 0.09. For uneven PPBs, >10 cells;Wilcoxon rank sum test p-value = 4.08E-03. For 1-sided PPB: n = 14 cells. For misoriented PPBs: n = 9 cells. D). *dcd3a-2/+ dcd3b/+* and E-G). *dcd3a-2 dcd3b*, representative images CFP-TUBULIN (green) and TAN1-YFP (magenta) time-lapse. Time indicated in hr:min. Scale bar = 10 µm.

**Table 1.**
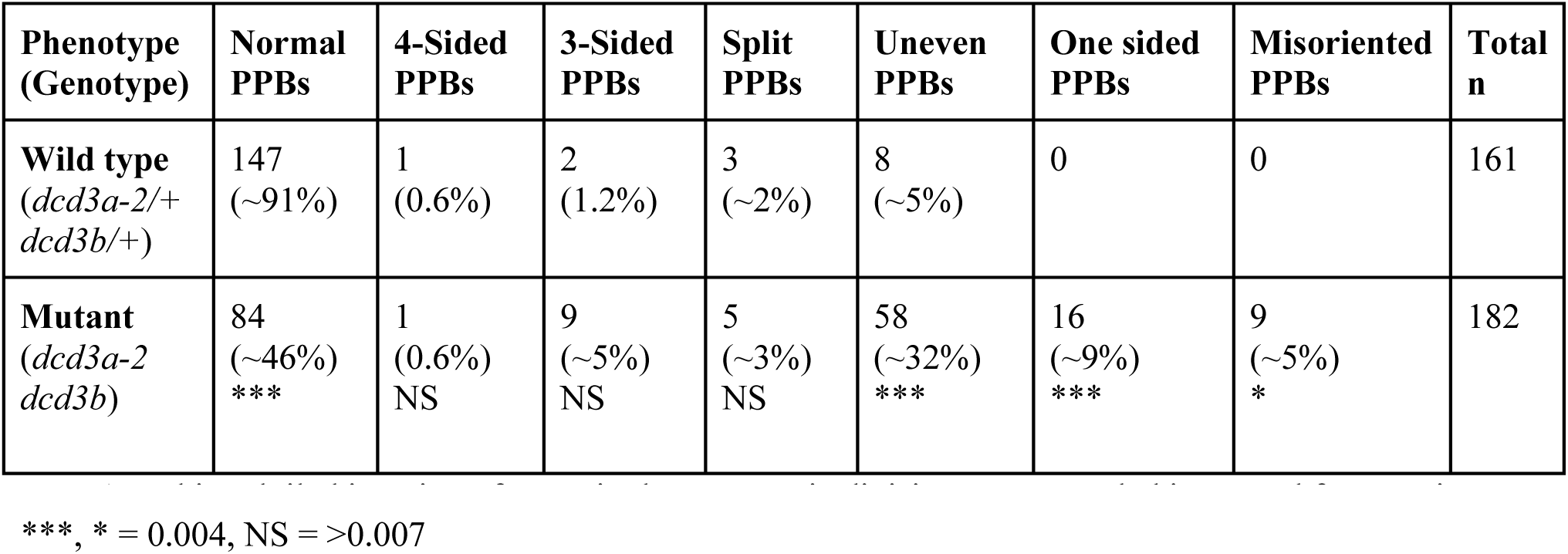
Unbiased tiled imaging of PPBs in the symmetric division zone revealed increased frequencies of abnormal PPBs in the *dcd3a-2 dcd3b* mutants compared to wild-type siblings. At least three 4 week-old plants per genotype were used. Fisher exact test (with Bonferroni correction of .05/7 = 0.007) < 0.001 ***, * = 0.004, NS = >0.007

| Phenotype (Genotype) | Normal PPBs | 4-Sided PPBs | 3-Sided PPBs | Split PPBs | Uneven PPBs | One sided PPBs | Misoriented PPBs | Total n |
| --- | --- | --- | --- | --- | --- | --- | --- | --- |
| **Wild type**<br>(*dcd3a-2/+*<br>*dcd3b/+*) | 147<br>(~91%) | 1<br>(0.6%) | 2<br>(1.2%) | 3<br>(~2%) | 8<br>(~5%) | 0 | 0 | 161 |
| **Mutant**<br>(*dcd3a-2*<br>*dcd3b*) | 84<br>(~46%)<br>*** | 1<br>(0.6%)<br>NS | 9<br>(~5%)<br>NS | 5<br>(~3%)<br>NS | 58<br>(~32%)<br>*** | 16<br>(~9%)<br>*** | 9<br>(~5%)<br>* | 182 |

To determine if one-sided or uneven PPBs alter the timing of *dcd3a-2 dcd3b* symmetric divisions, we re-analyzed the time-lapse imaging from Supplementary Table 2. Rare one-sided PPB cells tended to have both more variable and delayed division times (Supplementary Table 6). Those cells with uneven PPBs also had more variable timing, but overall division times were not significantly different than cells with normal PPBs.

Next, one-sided or uneven PPB division trajectories were analyzed to show that they do not alter division plane positioning in *dcd3a-2 dcd3b* mutants (Figure 5D-G, n > 3 plants). PPBs and division planes were positioned correctly in wild-type divisions (n = 69/69 cells). Similarly, *dcd3a-2 dcd3b* double mutant cells with normal (n = 17/17 cells) and uneven (n = 23/23 cells) PPBs divided in a normal position. One-sided PPBs also generated apparently properly oriented symmetric divisions that bisect the cell in half (n = 7/7 cells, Figure 5D-F). Together this indicates that in symmetrically dividing cells, final division plane positioning is not aberrant even when the PPB is only partially formed.

Misoriented PPBs in the *dcd3a-2 dcd3b* mutants generated similarly positioned misoriented divisions (n = 11 from 4 plants, Figure 5G). We were able to detect these rare divisions, by using cells in metaphase and telophase with TANGLED1-YFP at the cell cortex as a proxy for the former PPB location, as previously described^47^. This correlation of misoriented PPBs and division planes strongly suggests that the resulting aberrant cell shapes are due to misoriented PPBs, not due to phragmoplast guidance defects. Together, these results further suggest that the majority of division plane defects in symmetrically dividing cells in the *dcd3a-2 dcd3b* double mutant are due to misoriented PPB positioning.

### PPB abnormalities influence nuclear positioning

While analyzing images, we observed that the nucleus in *dcd3a-2 dcd3b* mutants was often aberrantly close to the PPB, particularly when the PPB was partially formed. We measured the distance between the nucleus centroid and the cell centroid, and found that, in interphase, there was no significant difference in the nuclear positioning between mutant and wild-type cells (Supplementary Figure 7; n > 54 cells from 3 plant of each genotype; Wilcoxon test p-value = 0.7). However, when cells were in late G2/prophase, nuclear centroids in *dcd3a-2 dcd3b* cells were often further away from the cell center compared to wild-type nuclear centroids (Supplementary Figure 7; n > 112 cells from 3 plants; Wilcoxon test p-value = 4.2E-07).

To determine whether the PPB influenced nuclear positioning, we obtained the normalized distance between the nucleus centroid and the PPB from micrographs of *dcd3a-2 dcd3b* and wild-type siblings (Figure 5A-C). Similar to wild-type sibling nuclei, the nuclei of *dcd3a-2 dcd3b* mutants with normal and 3-sided PPBs were close to the cell center (Figure 5A-C; p-value > 0.05). However, in cells with uneven, one-sided, or mispositioned PPBs, the nucleus was typically located closer to the more abundant microtubule accumulations in *dcd3a-2 dcd3b* cells (Figure 5A-B), with 2-fold or higher distance between nuclear and cell centroids (Figure 5C, Wilcoxon rank sum, p-value = 4E-03). Together, these data suggest that PPB position influences the positioning of the nucleus in *dcd3a-2 dcd3b* mutants.

## Discussion

Multiple independent analyses across several developmentally distinct leaves and roots with different growth conditions showed that *dcd3a-2 dcd3b* double mutants generated small, aberrantly-shaped organs. Further, leaf and root elongation rates were reduced. Small, less elongated cells are a hallmark of *p60* mutants in other plants^15,17,57,59^ but how cell shape defects contribute to overall growth has not been analyzed in detail. Here we have shown that both cell elongation and reduced frequency of cell division contribute to small organs, although cell elongation may play a dominant role.

Cell size thresholds contribute to the switch between G1 and S phases of the cell cycle^64^ and also between G2 and M ^63^. It is plausible that slower cell growth delays G1 and G2 progression in the *dcd3a-2 dcd3b* double mutant. We speculate that G1 or G2 delays may be the primary drivers for reductions in the number of actively dividing cells in the *dcd3a-2 dcd3b* mutant, since division itself was not delayed.

Observations from Arabidopsis shoot apical meristems show that additional time is typically required for small cells to complete the cell cycle^63^, despite smaller cells growing more quickly than larger neighboring cells^65^. Interestingly, blocking Arabidopsis meristem cells at or before S phase with the DNA synthesis inhibitor aphidicolin also prevents cell expansion, highlighting the interconnections between cell cycle progression and cell elongation^66^. In the future, it would be useful to use recently developed live-cell markers to directly measure G1-S-G2-M transitions and proportions^63,67^ to directly test the hypothesis that G1 and G2 are delayed.

Considerable variation exists in microtubule severing frequencies within different Arabidopsis cell types, varying up to five-fold in hypocotyl and tenfold in pavement cells^6,68^. We also observed variability in severing frequencies among elongating pavement cells from different wild-type maize plants (∼ 4-5 X 10^−3^ events/μm^2^/min), potentially reflecting differences in genetic background. In our study, *katanin p60* mutant combinations typically had ∼2-fold reduced microtubule severing frequencies compared to wild-type siblings. However, microtubule severing in maize epidermal cells was not completely eliminated, in contrast to microtubule severing frequencies of the *p60* mutant in the Arabidopsis petiole^20^, and cotyledons^6,69^. Similarly, lack of p80 subunits in the Arabidopsis stamen reduced severing frequency ∼11-fold^70^. One hypothesis for low but detectable levels of severing in mutants is that some allele combinations may reflect only partial loss of function. For example, *dcd3a-1*, the allele in the inbred CML228, contains many SNPs but no overtly deleterious amino acid substitution, and generates a transcript (Supplementary Figure 2D). Relative reductions in severing are smaller in *dcd3a-1 dcd3b* mutants compared to *dcd3a-2 dcd3b,* suggesting that some mutant combinations may reflect partial reduction in activity. An alternate hypothesis is that other microtubule severing proteins may contribute more to severing in some monocots. For example, spastin plays a significant role in rice plant growth^71^, while Fidgetin-like in maize has no described vegetative role^72^. Determining whether severing still occurs in maize *katanin* mutants due to contributions from other microtubule severing proteins or due to residual partial function of the mutants in maize awaits further study.

A significant fraction of PPBs in the *dcd3a dcd3b* double mutant were aberrantly formed, generating uneven PPBs and occasional frayed or, in rare cases, one-sided PPBs, similar to those observed in Arabidopsis roots and petioles^20,28,33^. Time-lapse analysis showed that failure to form a morphologically normal PPB led to division delays, similar to delays in division seen in Arabidopsis *katanin* mutants ^20^. The rarity of one-sided PPBs likely contributes to overall similarity in division times between mutant and wild-type cells.

The majority of PPBs, even those that are aberrantly formed, appear correctly localized and generate properly oriented divisions in symmetrically dividing cells of the *dcd3a-2 dcd3b* double mutant. That poorly formed or undetectable PPBs do not cause major division positioning defects has also been observed in the Arabidopsis *tonneau recruiting motif* (*trm678)* triple mutant, as well as in the *tonneau1a* single mutant, which have greatly compromised PPBs but variable division positioning defects depending on cell type^73,74^. In contrast, lack of PPB formation is tightly correlated with mispositioned division in asymmetric, subsidiary cell divisions^47^. Together, this suggests that the PPB may play differential roles in division positioning in different cell types, with its role in division positioning more clearly revealed in asymmetric divisions.

Rare misoriented PPBs in symmetrically dividing cells of the *dcd3a-2 dcd3b* double mutant generated aberrant small cells with curved cell walls. In Arabidopsis roots, ectopic and mispositioned longitudinal divisions within *katanin* mutants generated additional protoxylem and calyptrogen cells that form additional xylem and root cap cells respectively^33^. Additionally, *katanin* mutants also generated mispositioned hair and non-hair cells in roots. Interestingly, aberrant cell-type specification was seen in cell files with apparently normal division planes suggesting that, although the mutant has misoriented division plane positioning, that alone did not cause the misspecified cell fates^14^. Similar root hair and division mispositioning occurs in the *sabre* mutant^75,76^. Another fascinating division positioning defect generates a similar small and aberrantly shaped cell in ectopically dividing endodermal cells in the *inflorescence and root apices receptor kinase (irk)* mutant^77^. These small cells eventually transition into cortex cells. Whether the PPB is mispositioned in the *irk* mutant is not yet known.

Our examination of nuclear positioning defects highlights an important role of both the PPB, and KATANIN, in correctly orienting the nucleus during late G2/prophase. Connections between the PPB and the nuclear envelope have been observed in many organisms: these “bridge microtubules” are thought to promote bipolar spindle formation^29,78–81^. Interestingly, drugs that inhibit microtubule depolymerization disrupt mid-cell nuclear positioning in cells with PPBs indicating that dynamic microtubule organization is critical for nuclear positioning^82^. Here we showed that defects in PPB morphology were correlated with mispositioned nuclei: nuclei were located closer to more microtubule accumulation in *katanin* mutants. KATANIN-GFP accumulates at the nuclear envelope and the PPB during prophase/G2 with a proposed role in severing microtubules to establish bipolarity of the prophase spindle^28^. Here we suggest that it may also sever microtubule connections at the nuclear envelope to adjust nuclear position in relation to the PPB.

## Materials and Methods

### Plant Growth Conditions

To generate plants with specific genotypes, maize plants were grown to maturity and crossed in standard field conditions at UCR, University of North Texas, and Purdue University.

Maize plants were grown in the greenhouse with the following conditions: 1 L pots with soil (20% peat, 50% bark, 10% perlite, and 20% medium vermiculite) supplemented with magnesium nitrate (50 ppm N and 45 ppm Mg), calcium nitrate (75 ppm N and 90 ppm Ca), and Osmocote Classic 3-4M (NPK 14-14-14 %, AICL SKU#E90550). Plants were grown under standard greenhouse conditions (∼31-33°C with supplemental lighting from 5-9 PM at ∼400 µEm^-2^s^-1^). For analysis of the fully expanded fifth and eighth leaves, plants were grown for 5 weeks and 12 weeks, respectively. For time-lapse and other imaging, plants were grown for 4 weeks.

For first leaf and root analysis, germination paper was used to germinate kernels after sterilization according to ^83,84^ and grown for 10-11 days in a growth chamber (Percival) at 24°C with 16-h white light ∼111 µE*m−2s−1 (F17T8/TL741 Fluorescent Tube (Philips)) and 8-h dark cycles.

For leaf elongation rate experiments, maize kernels were germinated in 1 L pots as described above and grown in a growth chamber (Percival PGC-40L2) at constant temperature of 32 °C with 16-h light (∼400 µEm^-2^s^-1^) and 8-h dark cycles.

### Mapping *Clt1-1* to a lesion in an ortholog of AtKTN1 on Chr 8

*Clt1-1* was mapped to an interval on the long arm of chromosome 8^52^ that proved later to be ∼21.8 Mb in length. To refine the interval for *Clt1-1* further, the mutant, which was generated by Dr. Gerry Neuffer using ethyl methanesulfonate (EMS) in an unknown genetic background, was introgressed for eight generations into the B73 inbred maize background. We defined a 6.6Mb, non-B73 linkage block on Chr 8, flanked on both sides by homozygous B73 loci, containing *Clt1-1* (Supplemental Figure 1).

Genotyping 259 more individuals from a B73 X *Clt1-1*/+ (B73) cross identified a recombinant that reduced the Clt1-1 interval further with a new right flanking marker, ss230245925, at position 150,879,259 bp (Supplemental Figure 1). No polymorphic markers could be identified to demarcate the left boundary of the non-B73 linkage block; the closest markers (ss230245196 and ss230245225) were homozygous for the B73 allele. We conservatively estimated marker ss230245130 as the putative left boundary instead, leaving an interval containing 16 genes that might underlie *Clt1-1*.

To identify the causative lesion within these 16 genes, genomic DNA from two *Clt1-1* homozygotes in a B73 background were sequenced. After aligning the reads to the B73 genome sequence (AGPv3.21; 6.4X and 6.7X coverage, respectively) and filtering out low-quality variant calls, plotting the number of non-B73 sites in genomic windows along Chr 8 (Supplemental Figure 1) confirmed the interval obtained through positional cloning. The only mutation within the interval that affected the coding sequence of any gene was a single missense mutation in an AtKTN1 ortholog, ZEAMMB73_183479 (also known as GRMZM2G017305 at Chr 8:150,792,636-150,799,106 AGPv3; Supplementary Figure 1). The location of this gene coincided with a region of sharply reduced frequency of variant sites. The putative mutation had a Phred-scaled P-value from Fisher’s exact test for strand bias (FS) of 0 indicating no strand bias during sequencing. The mutation is a C to T transition, which would change a serine to phenylalanine in the ATPase domain of the protein (Figure 1A), predicted to be deleterious (SIFT score of 0). This SNP was confirmed by both a dCAPS marker and Sanger sequencing, which also showed that there is 100% identity with B73 at all other positions in this gene, including the 5’ untranslated region (UTR), exons, introns and the portion of the 3’ UTR that was sequenced. These results strongly suggest that mutation of ZEAMMB73_183479 is the cause of *Clt1-1*.

### Genotyping

DNA was extracted from leaves or kernels using standard protocols^85,86^. Mutant alleles (*dcd3a-1*, *dcd3a-2*, *dcd3b*, *Clt1*) and/or transgenes (YFP-ɑ-TUBULIN, CFP-β-TUBULIN, TAN1-YFP) were genotyped using 0.5 µM primers in Supplementary Table 7. Restriction enzyme digests followed PCR with SspI, MspI, and DraI for *dcd3a-1*, *dcd3b*, and *Clt1*, respectively. Presence of transgenes were also verified through painting leaves with 4 g/L glufosinate (Finale, Bayer) in 0.1% Tween 20 (Sigma), and scoring for resistance after 5-7 days.

### Confocal Microscopy

Images were acquired using a Yokogawa W1 spinning disk microscope on a Nikon Eclipse TE inverted stand with an EM-CCD camera (Hamamatsu 9100c) built by Solamere Technology Inc. Solid-state Obis lasers (40-100 mW) were used in combination with standard emission filters (Chroma Technology). For YFP-TUBULIN and TAN1-YFP, a 514 nm laser with emission filter 540/30 nm was used. For CFP-TUBULIN, a 445 nm laser with emission filter of 480/40 nm was used. A water immersion objective (60X/1.2 NA) and oil objective (100X/1.45 NA) were used. Images and time-lapses were taken with Micromanager-1.4 using a 3-axis DC servo motor controller and ASI Piezo Z stage. For microtubule severing time-lapse, 2 second time intervals were used. For cell division time-lapse, 10-minute time intervals were used. When Z-stacks were acquired, a Z-interval of 2 microns was used.

For microtubule severing, some images were collected using a Zeiss LSM 880 confocal laser scanning microscope (100X oil objective immersion lens, NA = 1.46) with Airyscan. A 514 nm laser with bandpass filters of 465-505 nm with a long-pass 525 filter was used with a time interval of 2 seconds. Images were processed using default Airyscan settings with Zen software (Zeiss).

### Microtubule Severing

Four- to five-week-old plants expressing YFP-ɑ-TUBULIN or CFP-β-TUBULIN were dissected to the youngest emerging leaf. Tissue samples were obtained from the elongation zone (7 cm from the ligule for wild-type siblings and 4 cm for *dcd3a-2 dcd3b*) and mounted on water in a rose chamber as described^87^. Samples were imaged every 2 seconds. Time-lapses were bleach corrected (Simple Ratio) and re-aligned using the Translation selection in StackReg in FIJI^88^. Only cells with similar microtubule density and anisotropy were analyzed. Individual cells were selected with the Rotated Rectangle Tool and cell area was measured. Microtubule density and anisotropy were calculated for the first time frame of bleach corrected (Simple Ratio) images using BoneJ^89^ and FibrilTool^90^ plugins, respectively, in FIJI. For microtubule density, images were thresholded using default segmentation prior to obtaining the Area/Volume Fraction using the BoneJ plugin. Microtubule density was calculated as 1 - Area/Volume. For microtubule severing analysis, a microtubule severing event was identified by depolymerization of one or more microtubules at a microtubule crossover.

### Root and leaf growth experiments

Maize kernels were sterilized according to ^91,92^, with the following modifications. Maize seeds were incubated in 80% ethanol for 3 minutes, rinsed with sterile water, incubated with 50% Bleach for 15 min, and rinsed with sterile water. Kernels were imbibed overnight in sterile water with rocking. Growth assays were conducted following the published method^84^. Sterilized maize seeds germinated on germination paper (Fisher Scientific #NC1466201) soaked in 2.5 g/L of Captan 50 fungicide (Southern Ag #01600), rolled and stored vertically in beakers filled with 400 mL of 0.5X pH5.7 Linsmaier and Skoog media (Caisson Labs #LSP03-50LT). The media was changed every 2 days. Root images with a 30.5 cm ruler were taken every 24 hours for 10 days with a Canon PowerShot SX540 HS digital camera. Germination was determined by the protrusion of the radicle from the coleorhiza and root measurements excluded the coleorhiza. Root lengths were measured manually using FIJI.

Maize kernel DNA was genotyped^86^ and two wild-type and two mutant kernels were planted in the growth chamber every day for 10 days. Upon leaf 5 emergence, leaf 5 length was measured using a 30.5 cm ruler every 24 hours for 7 days to obtain the leaf elongation rate^93^.

### Leaf area and cell measurements

Fully expanded leaf 5 and 8 were obtained from at least 3 *dcd3a-2/+ dcd3b/+* and *dcd3a-2 dcd3b* mutant plants grown in the greenhouse. Leaf 1 was obtained from seedlings grown in germination paper rolls for 11 days. Whole leaf images were taken next to a 30.5 cm ruler using a camera (Canon powershot SX540 HS). Glue impressions^94^ were made from the base, middle, and tip regions of the leaves and imaged with a Nikon light compound microscope using a 4X or 10X objective with a 2X AmScope camera attachment. Leaf areas, pavement cells, and stomatal complexes were outlined manually in FIJI.

### Approximating Total Cell Counts in Leaves 1, 5, and 8

The number of cells in a defined area was used to approximate the number of cells in the whole leaf. The defined areas consisted of at least 3 micrographs each from the tip, middle and base of the leaf. Averages for each section were calculated separately and then combined as follows. PC = total pavement cells measured; SC = total stomatal complexes measured; PCA = pavement cell area; SCA = stomatal complex area.

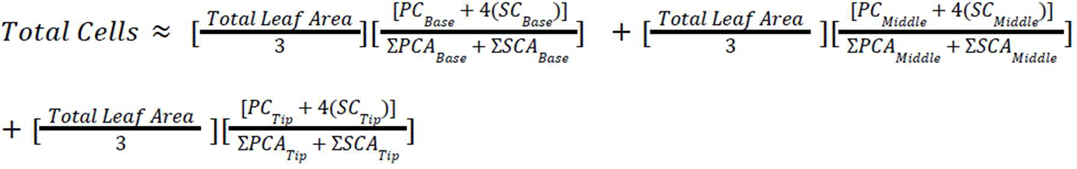

### Leaf Projections

**Projection 1** was calculated by multiplying the average cell length and width of *dcd3a-2 dcd3b* mutants by the estimated number of cells along the long and short axis of each *dcd3a-2/+ dcd3b/+* leaf.

**Projection 2** multiplied the average cell area of *dcd3a-2 dcd3b* mutants by the estimated cell number for *dcd3a-2/+ dcd3b/+* leaves. **Projection 3** multiplied the average estimated cell number of *dcd3a-2 dcd3b* mutants by the cell dimensions of each *dcd3a-2/+ dcd3b/+* leaf. **Projection 4** multiplied the average estimated cell number by the average cell area of *dcd3a-2 dcd3b* mutants. All projections were made to scale in GIMP.

### Cell Division and Nuclear Positioning Analysis

4-week old plants were used to obtain a leaf with a ligule height < 2 mm. Samples were taken within 0.5 cm from the ligule of the emerging leaf. The abaxial side was mounted in water in a rose chamber.

Time-lapse of symmetric cell divisions of plants expressing either YFP-TUBULIN, or CFP-TUBULIN and TAN1-YFP were obtained in 10 minute intervals. For metaphase time, the duration of the first frame without a PPB to the last frame before anaphase was counted. For telophase time, the duration of the first frame with a phragmoplast to the frame up to phragmoplast disassembly was counted. For phragmoplast expansion time, the first frame with a phragmoplast to the frame before the phragmoplast reached the cortex of the cell was counted.

### Figure Preparation

Figures were made using Gnu Image Manipulation Program (v2.10.38, https://www.gimp.org/) or Powerpoint for Supplementary Figures 1 and 2. Levels were adjusted linearly. If applicable, images were enlarged or rotated with no interpolation.

### Graphs and Statistics

Graphs and statistics were generated using R and RStudio using the following packages: tidyr^95^, ggplot2^96^, ggprism^97^, paletter^98^, and stringr^99^.

## Data availability

Raw reads for *Clt1-1* homozygotes have been deposited at NCBI SRA https://www.ncbi.nlm.nih.gov/bioproject/PRJNA344581.

## Acknowledgments

Thanks to Dr. Gerry Neuffer (University of Missouri) and the Maize Genetics Cooperation for generating and maintaining the *Clt1* mutant respectively. Thanks to Drs. Laurie Smith and Kim Gallagher for generating the *dcd3* mutant and Dr. Nicholas Miles for propagation. Thanks to Dr. George Peeters (Solamere Technology Inc) for troubleshooting the microscope issues. This work was supported by NSF grants (CAREER-IOS-1350874 AJW, CAREER-MCB-1942734 CGR, MCB-2426623 CGR, DBI-0822495 KL and CW, DBI-1922642 SEM, REU-2051131 CH, REU-2447384 ID), a training grant to Purdue University Agronomy from USDA-NIFA and the Purdue Research Foundation to KL and CW, undergraduate research funding from UCR-RISE to CI, CH, IC.

## Author contributions

Conceived and designed the analysis: AJW, CGR, CW, SEM, KL

Collected the data: SEM, KL, NM, LAA, CI, CH, ID, IC, PB, AJW, CW, CGR

Contributed data or analysis tools:

Performed the analysis: SEM, KL, CGR, AJW Wrote the paper: CGR, SEM, KL, CW, AJW

Revised and reviewed the paper: All authors reviewed the paper

Funding acquisition: SEM, CI, CH, ID, IC, AJW, CW, CGR

---

**Supplementary Figure 1.**
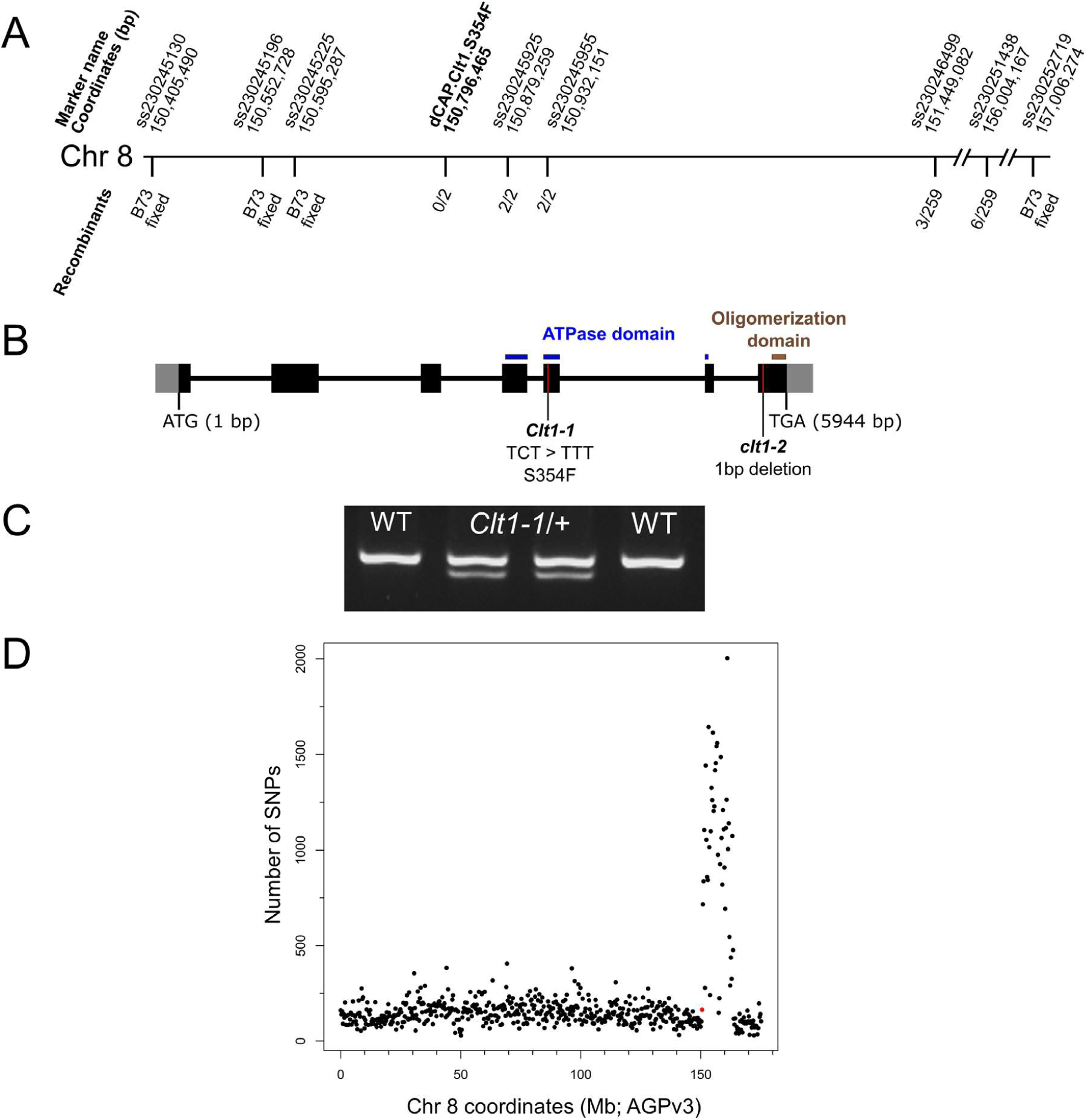
Mapping *Clt1* to the maize katanin ortholog *Dcd3b* on Chromosome 8. (A) Physical locations (B73 AGPv3) of markers used to genotype individuals from a *B73 X Clt1/+* mapping population and the number of recombinants observed at each marker. Where the number of individuals tested was only 2 instead of 259, only the plants recombinant for the outer markers were tested. (B) Gene model for *Dcd3b* indicating locations of *Clt1-1* and *dcd3b*/*clt1-2*. (C) A dCAP assay directly interrogating the putative causative C to T lesion in the *Dcd3b* locus. (D) Density of SNPs called from whole-genome sequencing of *Clt1* homozygotes, in 300 kb non-overlapping windows, with *Clt1* indicated in red.

**Supplementary Figure 2.**
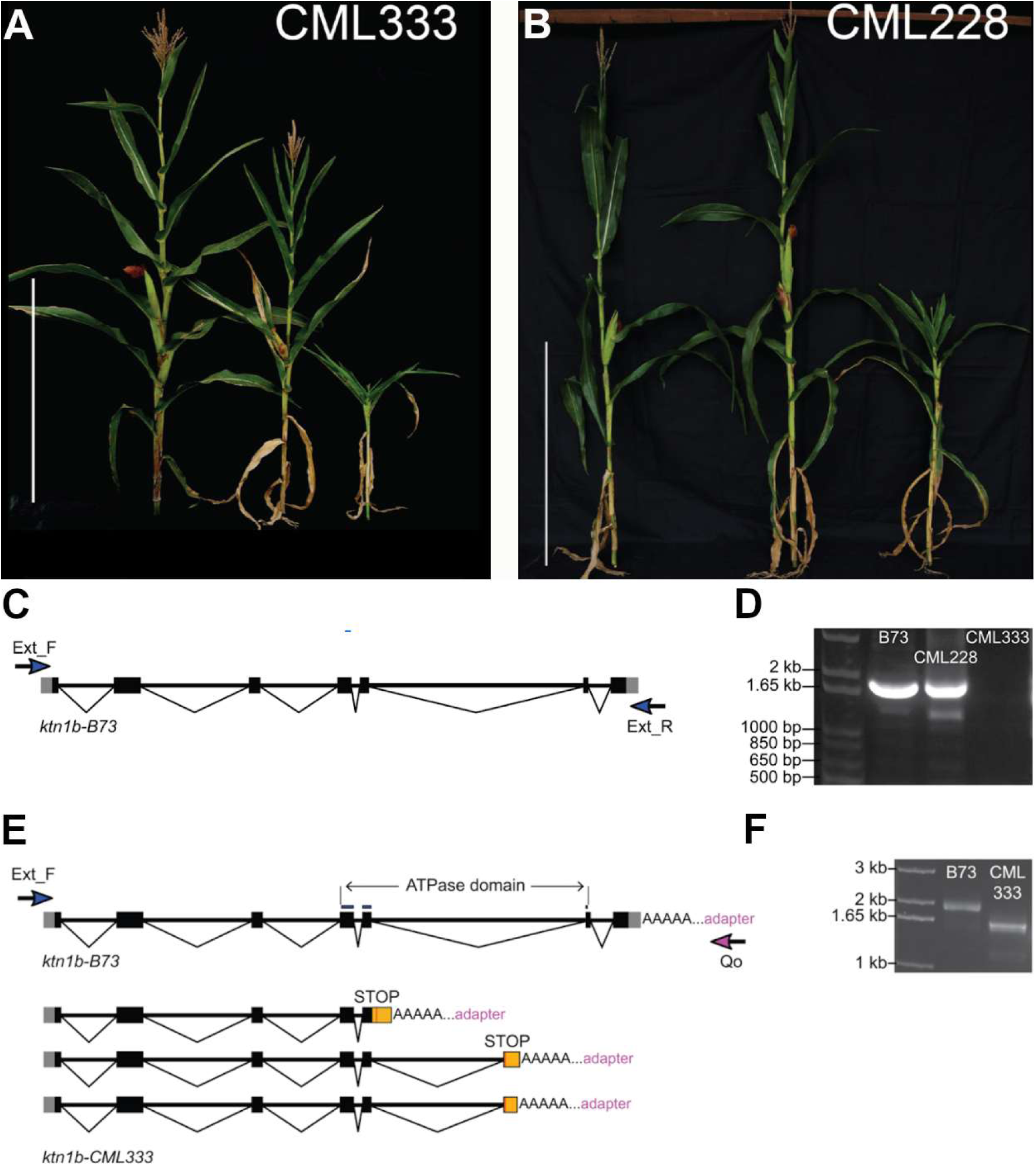
Two genetic enhancers of *Clt1/+* A) *Clt1/+ Dcd3a/Dcd3a* from B73 (left), middle is *Clt1/+ Dcd3a*/*dcd3a-2* (*dcd3a-2* from CML333), *Clt1/+ dcd3a-2/dcd3a-2* (right) B) *Clt1/+ Dcd3a/Dcd3a* from B73 (left), middle is *Clt1/+ Dcd3a*/*dcd3a-1* with *dcd3a-1* from CML228, (right) *Clt1/+ dcd3a-1/dcd3a-1*. Bar in each panel is 100 cm. C) Gene model of *dcd3a*/*ktn1b* in B73. Blue arrows indicate primers binding to the 5’ and 3’ UTRs and used to amplify the products shown in panel (D). (D) Amplification of *dcd3a/ktn1b* cDNA from CML228 and CML333 using the primers indicated in panel (C). Purified gel slices were cloned and sequenced. (E) Alternative splice forms of *dcd3a/ktn1b* transcript in CML333 revealed by sequencing of PCR products resulting from increasing PCR cycles from panel (D) and from (F) 3’ RACE. The earliest in-frame stop codons in the retained portions of intron 5 are indicated. (F) Amplification of CML333 *dcd3a/ktn1b* cDNA from 3’ RACE using the primers indicated in panel (E).

**Supplementary Figure 3.**
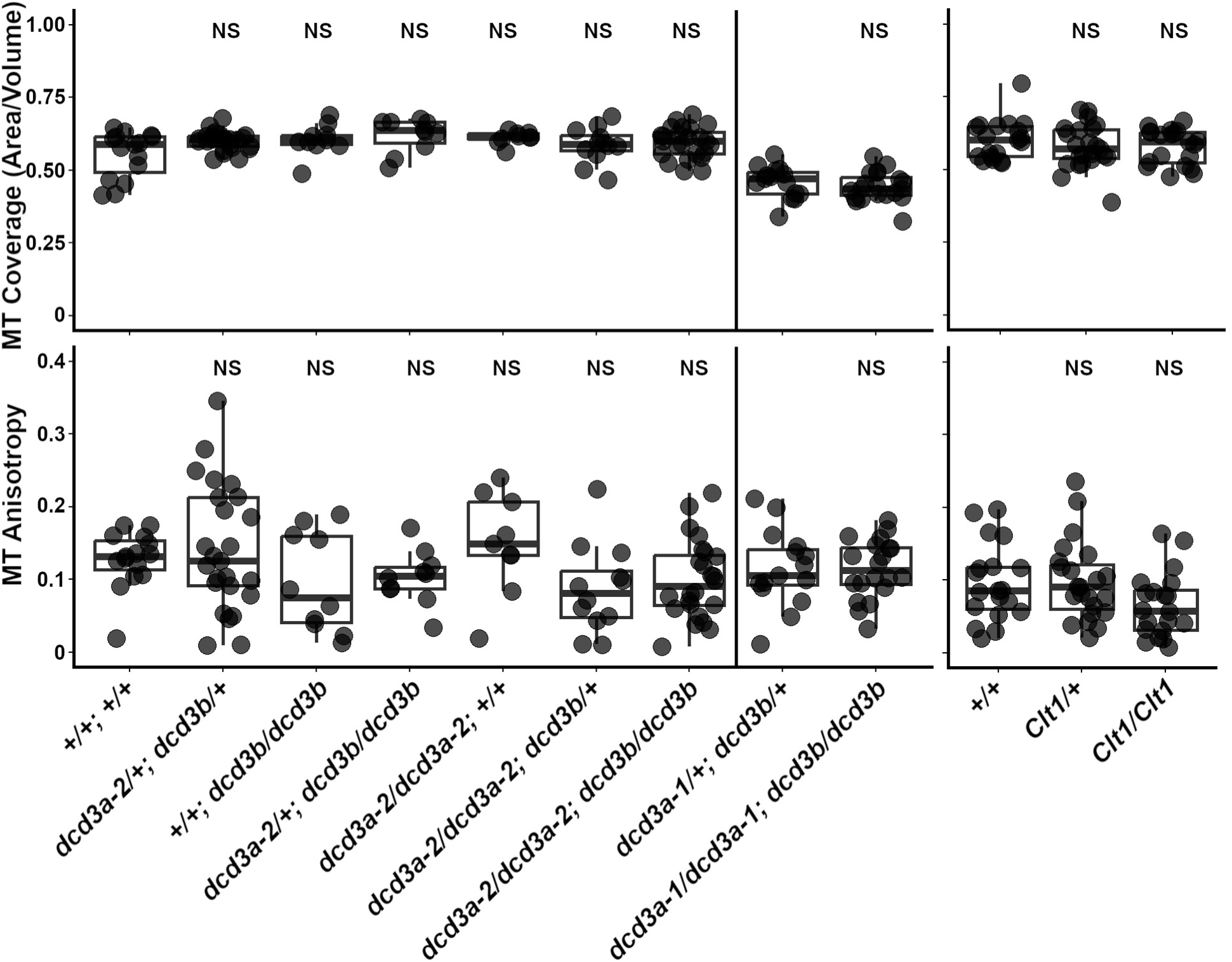
Microtubule coverage and anisotropy of cells selected for measuring severing in the elongation zone of the emerging leaf are not significantly different. Microtubule coverage was measured using the BoneJ plugin (see Materials and Methods). The average microtubule coverage for wild type (*+/+ +/+*) (0.554 ± 0.021 SE), *dcd3a-2/+ dcd3b/+* (0.601 ± 0.006 SE; p-value = 0.008), and *dcd3a-2 dcd3b* (0.593 ± 0.011 SE; p-value = 0.02). The average microtubule coverage for *dcd3a-1/+ dcd3b/+* and *dcd3a-1 dcd3b* was 0.460 ± 0.014 SE and 0.444 ± 0.011 SE, respectively (p-value = 0.374). The average microtubule coverage for wild-type (+/+) *clt1/clt1* (0.601 ± 0.016 SE), *Clt1/+* (0.579 ± 0.016 SE; p-value = 0.29), and *Clt1/Clt1* (0.580 ± 0.014 SE; p-value = 0.34). Microtubule anisotropy was obtained using FibrilTool (see Materials and Methods). The average microtubule anisotropy for *+/+; +/+* (0.128 ± 0.010 SE), *dcd3a-2/+; dcd3b/+* (0.142 ± 0.017 SE; p-value = 0.45), and *dcd3a-2 dcd3b* (0.099 ± 0.011 SE; p-value = 0.16). The average microtubule anisotropy for *dcd3a-1/+ dcd3b/+* and *dcd3a-1 dcd3b* was 0.115 ± 0.014 SE and 0.142 ± 0.009 SE, respectively (p-value = 1). The average microtubule anisotropy for +/+ (*clt1/clt1*) (0.097 ± 0.012 SE), *Clt1/+* (0.098 ± 0.011 SE; p-value = 1), and *Clt1/Clt1* (0.66 ± 0.010 SE; p-value = 0.06). P-values were obtained from pairwise t-tests (with Bonferroni correction .05/6=0.008 or .05/2=0.025).

**Supplementary Figure 4.**
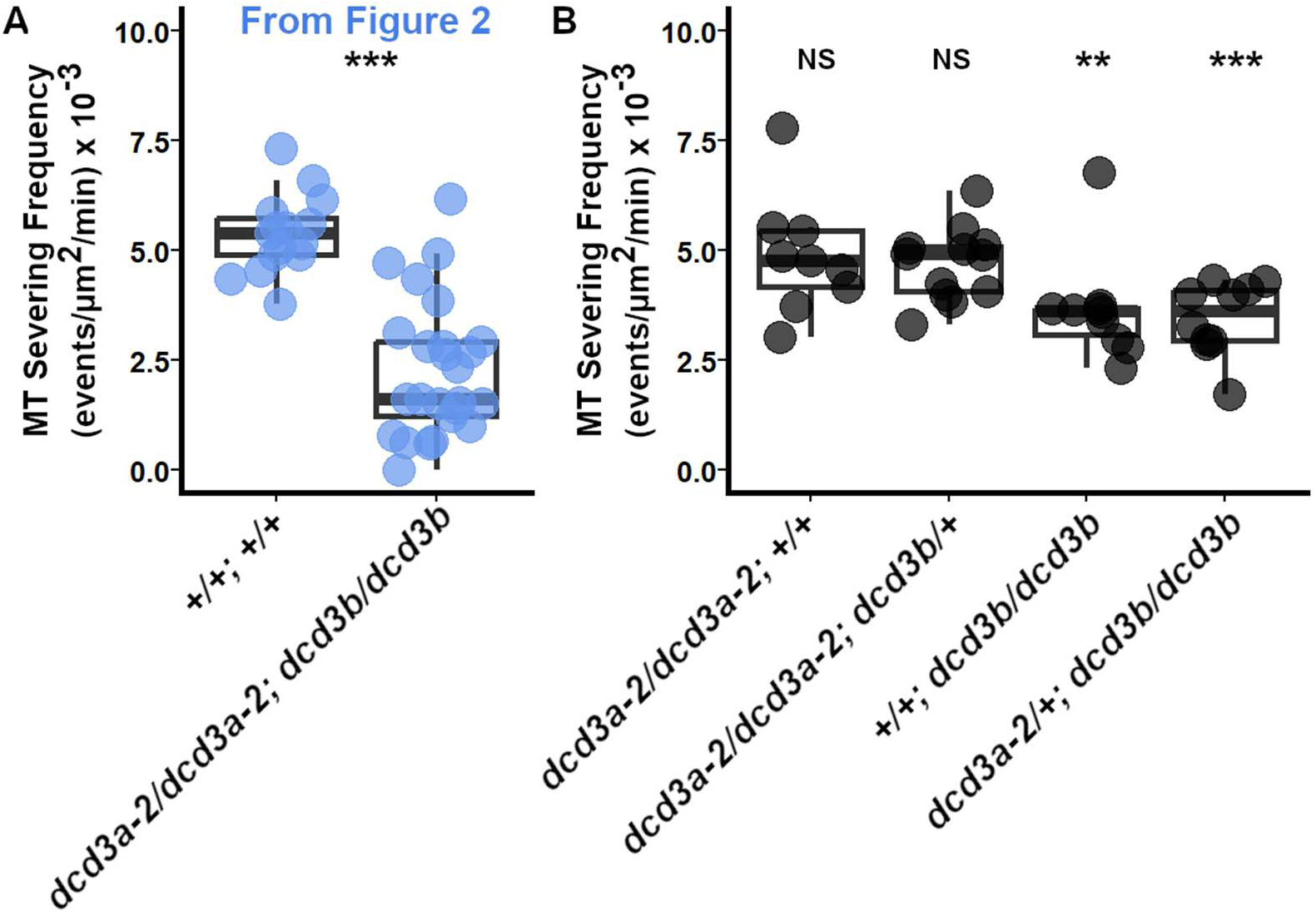
Microtubule severing frequency (events/µ^2^/min) X 10^-3^ in additional *katanin* mutant allele combinations. A). Microtubule severing frequency for +/+; +/+ and *dcd3a-2/dcd3a-2 dcd3b/dcd3b* taken from Figure 2. Pairwise comparison (with Bonferroni correction .05/4=0.0125) was done between *+/+; +/+* and other alleles shown in B. B). Microtubule severing frequency in *dcd3a-2/dcd3a-2; +/+* averaged 4.9E-03 events/µm^2^/min ± 0.04E-03 from n = 9 cells from 3 plants, p = 0.25; *dcd3a-2/dcd3a-2; +/dcd3b* averaged 4.7E-03 events/µm^2^/min ± 0.02E-03 SE from n = 12 cells from 6 plants, p = 0.09; *+/+; dcd3b/dcd3b* averaged 3.7E-03 events/µm^2^/min ± 0.04E-03 SE from n = 10 cells from 4 plants, p < 0.0001; *+/dcd3a-2, dcd3b/dcd3b* average 3.5E-03 events/µm^2^/min ± 0.02E-03 SE from n = 10 cells from 3 plants, p = 0.00002.

**Supplementary Figure 5.**
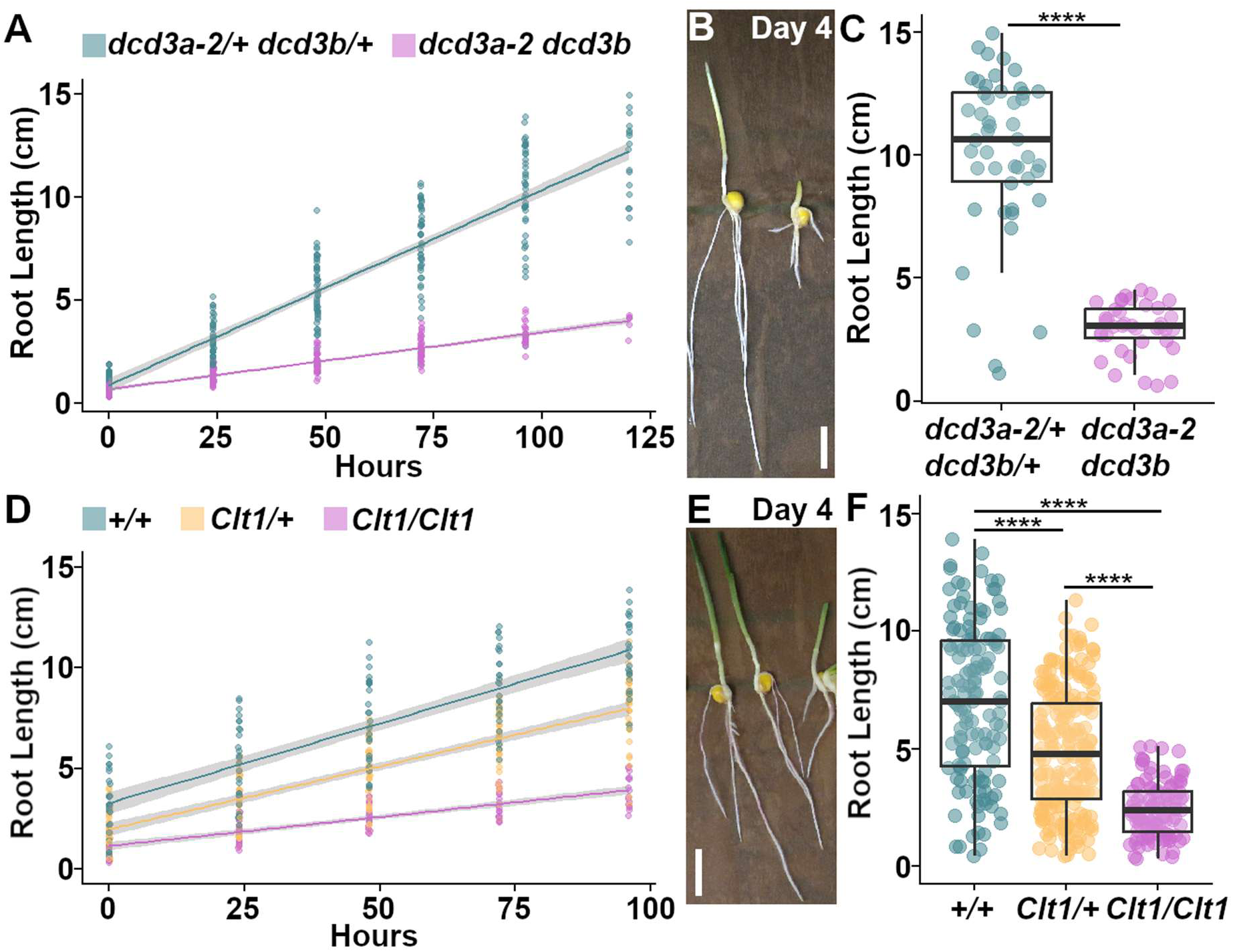
The *katanin* mutants have decreased root growth over time. (A). For the first five days after germination, *katanin* mutants had decreased linear root growth rate when compared to wild-type siblings (*dcd3a/+ dcd3b/+*: 2.4cm/day ± 0.07 cm/day; *dcd3a-2 dcd3b*: 0.48 cm/day ± 0.02 cm/day). (B). Representative *dcd3a-2/+ dcd3b/+* (left) and *dcd3a-2 dcd3b* (right) seedlings 4 days after germination. Scale bar = 2 cm. (C). Root length of *dcd3a-2/+ dcd3b/+* (10.3 cm ± 3.1 cm; n = 46 seedlings) and *dcd3a-2 dcd3b* (3.0 cm ± 1.0 cm; n = 39 seedlings) at 4 days post germination. Welch two sample t-test p-value < 2.2e-16. (D). Linear root growth rate for wild type (1.8 cm/day ± 0.2 cm), *Clt1*/+ (1.5 cm/day ± 0.1 cm), and *Clt1/Clt1* (0.7 cm/day ± 0.1 cm). (E). Representative wild-type (left), *Clt1/+* (middle), and *Clt1/Clt1* (right) seedlings at 4 days post germination. Scale bar = 2 cm. (F). Root length of wild type (10.3 cm ± 2.3 cm; n = 27 plants), *Clt1/+* (7.6 cm ± 2.2 cm; n = 44 plants), and *Clt1/Clt1* (3.6cm ± 1.1 cm; n = 22 plants) at 4 days after germination. **** asterisk indicates p-value ≤ 4e-12 from a pairwise t-test with Bonferroni correction.

**Supplementary Figure 6.**
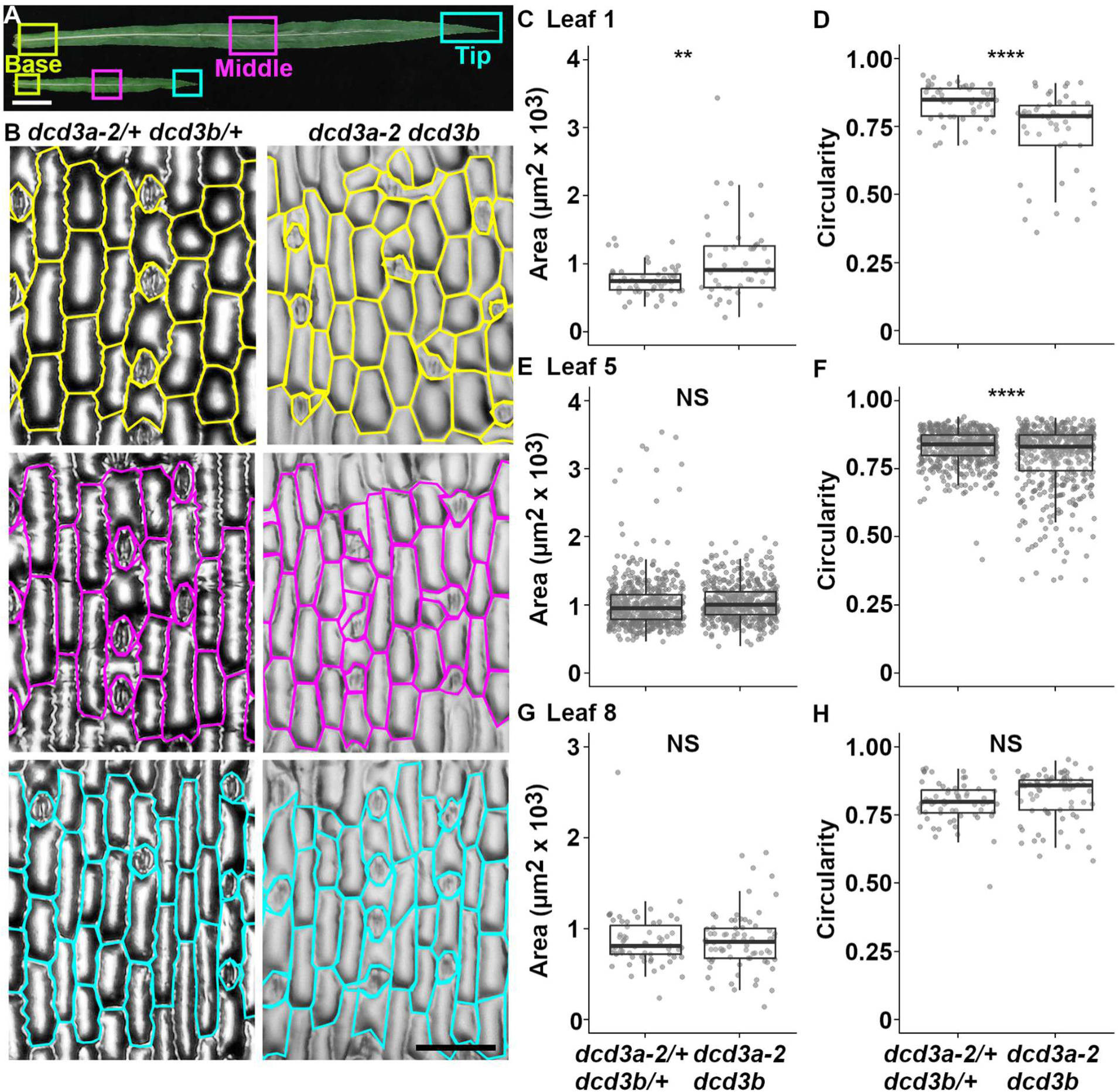
Area and circularity of stomatal complexes in *dcd3a-2 dcd3b* double mutants and wild-type siblings. (A) Base (yellow), middle (magenta), and tip (cyan) regions of leaf 1, 5 (shown here), and 8 were taken for glue impressions. Scale bar = 5 cm. (B) Representative examples of pavement cell and stomatal complex outlines from *dcd3a-2/+ dcd3b/+* and *dcd3a-2 dcd3b* glue impressions from the base (yellow outlined), middle (magenta outlines), and tip (cyan outlines) regions of the leaf. All images were taken at the same magnification. Scale bar = 100 µm. (C) Stomatal complex areas from leaf 1. (D) Circularity of stomatal complexes from leaf 1. (E) Stomatal complex areas from leaf 5 (F) Circularity of stomatal complexes from leaf 5. (G) Stomatal complex areas from leaf 8 (H). Circularity of stomatal complexes from leaf 8. Supporting data in Supplementary Table 2. Welch’s two sample t-test p-value < .01=**; < .00005= ****

**Supplementary Figure 7.**
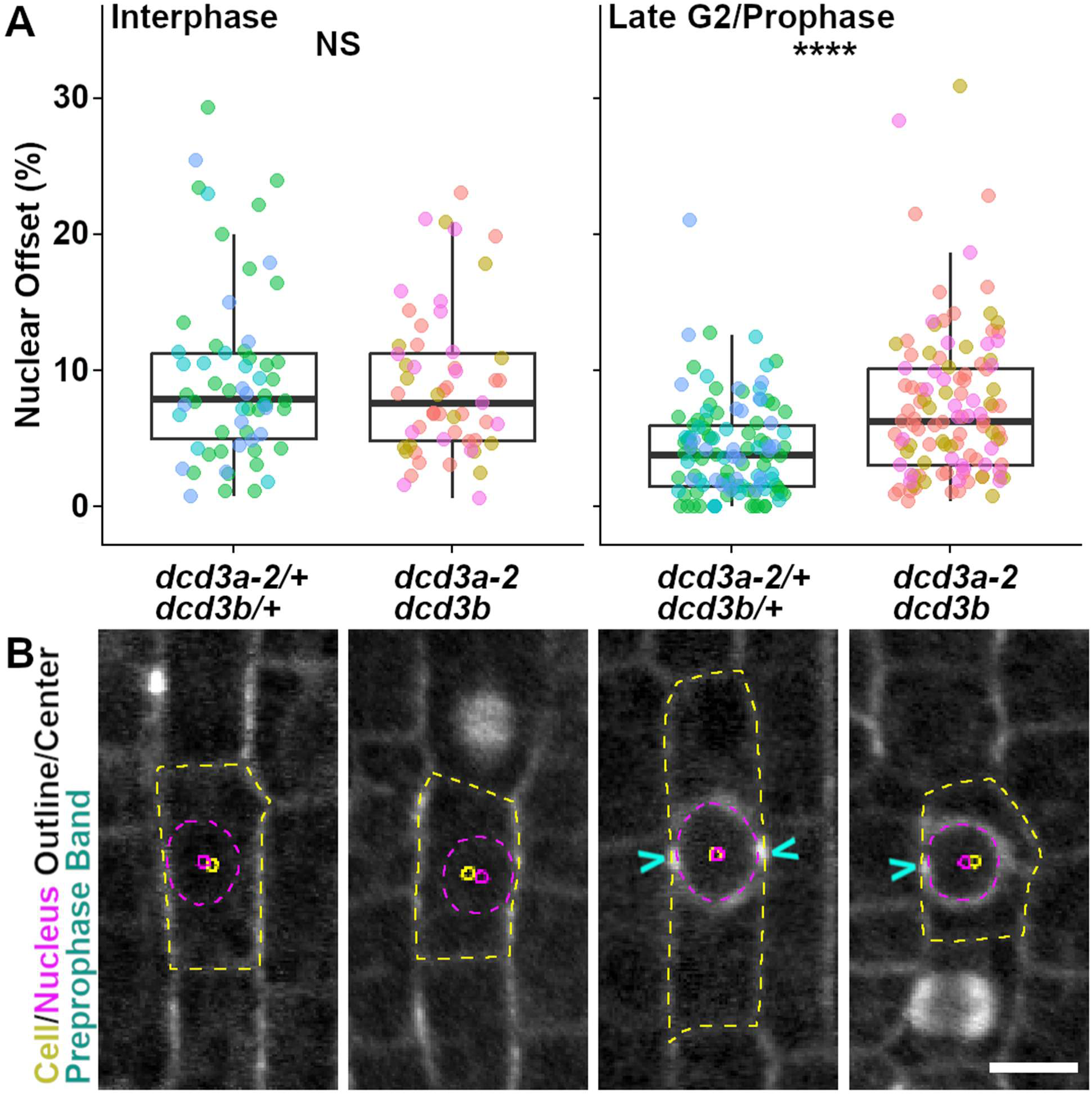
*dcd3a-2dcd3b* mutants have more offset nuclei during prophase when compared to wild-type siblings. (A). Boxplots showing normalized nuclear offset (%) from the cell center for cells in interphase and G2/prophase. For interphase: *dcd3a-2/+ dcd3b/+*: 9.5 ± 0.8, n = 62cells from 3 plants; *dcd3a-2 dcd3b*: 8.9 ± 0.7, n = 55cells from 3 plants. For G2/prophase: *dcd3a-2/+ dcd3b/+*: 4.2 ± 0.3, n = 117cells from 3 plants; *dcd3a-2 dcd3b*: 7.4 ± 0.5, n = 113 cells from 3 plants. Different colored dots = different plants. (B). Representative micrographs of *dcd3a-2/+ dcd3b/+* and *dcd3a-2 dcd3b* during interphase and G2/prophase. The cell (yellow) and nucleus (magenta) outline with their respective centroid is shown. The preprophase band is marked by the cyan arrows. Scale bar = 10 µm. Wilcoxon test p-value = 4.2E-07 = ****, Wilcoxon test p-value = 0.7 = NS

**Supplementary Table 1.** The *dcd3a-2 dcd3b* double mutants have smaller leaves with fewer cells than wild-type siblings. Leaf Area (cm^2^) ± standard error is shown. Approximate total cell count ± standard error is shown. N ≥ 3 plants per genotype per leaf. P-value from a two-tailed Student’s t-test P < 0.05 = *, P <0.005 = **.

| Leaf Number | Phenotype (Genotype) | Leaf Area (cm<sup>2</sup>) ± SE | P-value for Leaf Area | Total Approximate Cells ± SE | P-value for Total Cells |
| --- | --- | --- | --- | --- | --- |
| 1 | Wild type (*dcd3a-2/+ dcd3b/+*) | 5.7 ± 0.3 | 0.0002 ** | 4.2E+05 ± 3.4E+04 | 0.0020 ** |
|  | Mutant (*dcd3a-2 dcd3b*) | 1.6 ± 0.2 |  | 1.3E+05 ± 1.1E+04 |  |
| 5 | Wild type (*dcd3a-2/+ dcd3b/+*) | 145 ± 19 | 0.0060 * | 8.6E+06 ± 1.2E+06 | 0.0345 * |
|  | Mutant (*dcd3a-2 dcd3b*) | 51 ± 8 |  | 4.2E+06 ± 7.7E+05 |  |
| 8 | Wild type (*dcd3a-2/+ dcd3b/+*) | 473 ± 44 | 0.0147 * | 3.9E+07 ± 6.0E+06 | 0.0180 ** |
|  | Mutant (*dcd3a-2 dcd3b*) | 148 ± 9 |  | 1.4E+07 ± 7.3E+05 |  |

**Supplementary Table 2.** The *dcd3a-2 dcd3b* double mutants have smaller cells. Pavement cell and stomatal complex measurements for *dcd3a-2/+ dcd3b/+* and *dcd3a-2 dcd3b* leaves 1, 5, and 8. Welch’s two sample t-test p-value: NS > .05, ** < 0.01, *** < .005, **** < .00005. From 3 or 4 plants of each genotype. These data are used in Figure 3 and Supplementary Figure 6.

| Leaf | Phenotype (Genotype) | Total Pavement Cells | Median Pavement Cell Area ± SE | Median Pavement Cell Height ± SE | Median Pavement Cell Width ± SE | Total Stomatal Complexes Measured | Median Stomatal Complex Area ± SE | Median Stomatal Complex Circularity ± SE |
| --- | --- | --- | --- | --- | --- | --- | --- | --- |
| 1 | **Wild type** (*dcd3a-2/+ dcd3b/+*) | 153 | 2,781 ± 112 | 12 ± 3.6 | 23.7 ± 0.5 | 51 | 745 ± 31.7 | 0.85 ± 0.01 |
| | **Mutant** (*dcd3a-2 dcd3b*) | 404 | 1,557 ± 46 *** | 64 ± 1.5 **** | 24.7 ± 0.4 NS | 46 | 915 ± 88.3 ** | 0.79 ± 0.02 **** |
| 5 | **Wild type** (*dcd3a-2/+ dcd3b/+*) | 1,736 | 2,957 ± 31 | 113 ± 0.9 | 26.6 ± 0.2 | 496 | 953 ± 18 | 0.84 ± 0.003 |
| | **Mutant** (*dcd3a-2 dcd3b*) | 2,117 | 2,099 ± 17 **** | 80 ± 0.6 **** | 25.9 ± 0.1 **** | 465 | 1,005 ± 13 NS | 0.83 ± 0.005 **** |
| 8 | **Wild type** (*dcd3a-2/+ dcd3b/+*) | 204 | 2,554 ± 61 | 94 ± 2.3 | 26 ± 0.5 | 64 | 811 ± 39 | 0.80 ± 0.01 |
| | **Mutant** (*dcd3a-2 dcd3b*) | 336 | 1,719 ± 40 **** | 68 ± 1.3 **** | 26 ± 0.4 NS | 72 | 858 ± 38 NS | 0.86 ± 0.01 NS |

**Supplementary Table 3.**
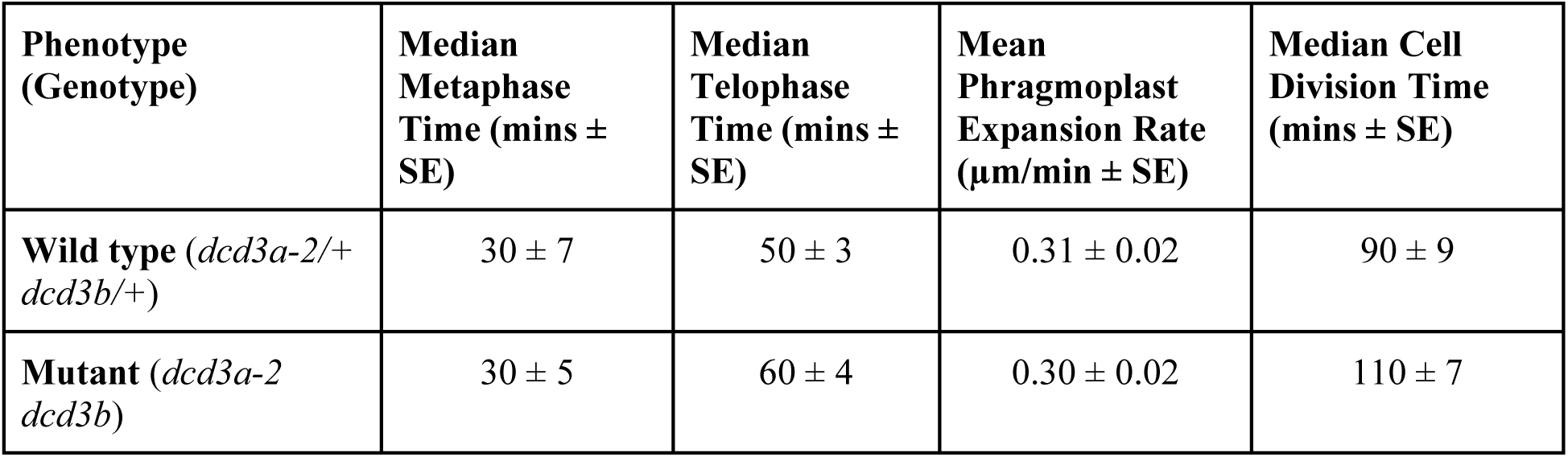
The *dcd3a-2 dcd3b* double mutants have similar cell division times as wild-type siblings. Data was obtained from n = 69 cells from 6 *dcd3a-2/+ dcd3b/+* plants and n = 48 cells from 8 *dcd3a-2 dcd3b* plants. Wilcoxon rank sum test p-values were > 0.05 for all measurements.

| Phenotype (Genotype) | Median Metaphase Time (mins ± SE) | Median Telophase Time (mins ± SE) | Mean Phragmoplast Expansion Rate (μm/min ± SE) | Median Cell Division Time (mins ± SE) |
| --- | --- | --- | --- | --- |
| **Wild type** (*dcd3a-2/+ dcd3b/+*) | 30 ± 7 | 50 ± 3 | 0.31 ± 0.02 | 90 ± 9 |
| **Mutant** (*dcd3a-2 dcd3b*) | 30 ± 5 | 60 ± 4 | 0.30 ± 0.02 | 110 ± 7 |

**Supplementary Table 4.** The *dcd3a-2 dcd3b* double mutants have similar proportions of late G2 and mitotic cells compared to wild-type siblings. . Data was obtained from at least three plants of each genotype for wild type plants (*dcd3a-2/+ dcd3b/+*) and mutant plants (*dcd3a-2 dcd3b*) . Fisher’s exact test p-values: NS: p> 0.05, * < 0.05, ** < 0.005 ***<0.00001. Data for Figure 4B. This data is a subset of the data from Supplementary Table 5.

| Mitotic Stage | Wild-type (*dcd3a-2/+ dcd3b/+*) | Mutant (*dcd3a-2 dcd3b*) |
| --- | --- | --- |
| Preprophase/prophase | 54% (101) | 59% (114) NS |
| Metaphase/anaphase | 12.8% (24) | 9.8% (19) NS |
| Telophase/cytokinesis | 33.2% (62) | 31.1% (60) NS |
| Total mitotic cells | 187 | 193 |

**Supplementary Table 5:**
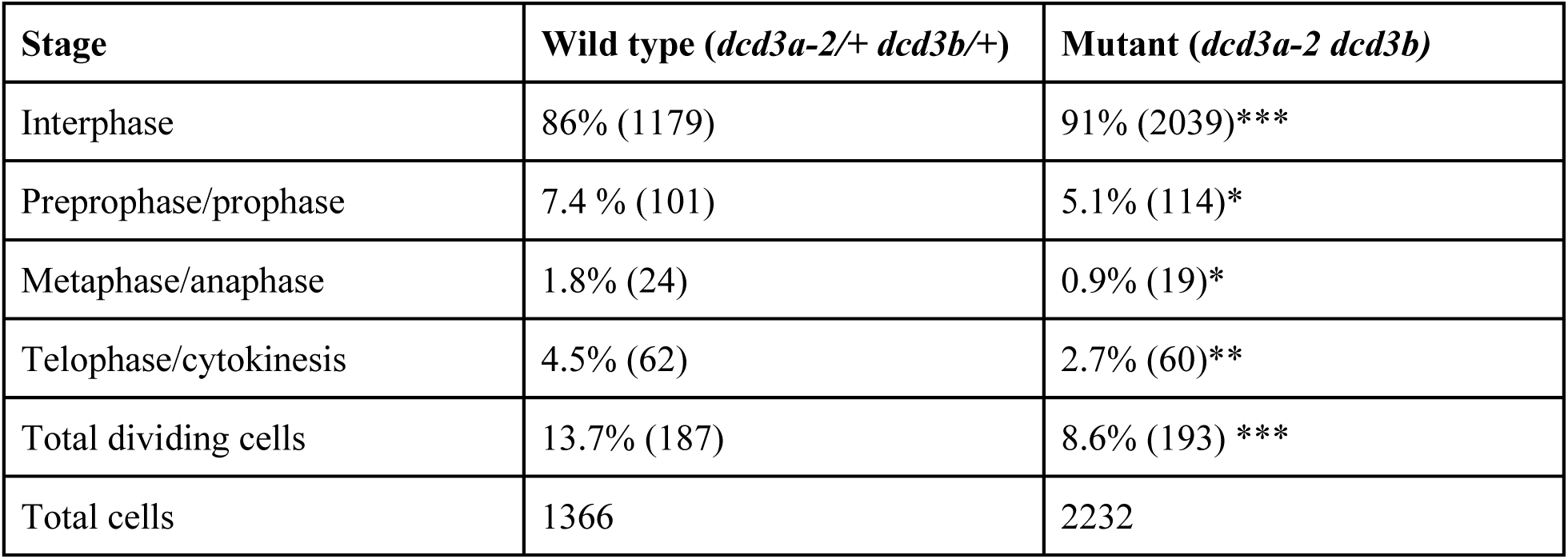
The *dcd3a-2 dcd3b* double mutants have less dividing cells compared to wild-type siblings. . Data was obtained from at least three plants of each genotype for wild type plants (*dcd3a-2/+ dcd3b/+*) and mutant plants (*dcd3a-2 dcd3b*) . Fisher’s exact test p-values: NS: p> 0.05, * < 0.05, ** < 0.005 ***<0.00001. Data for Figure 4B and Supplementary Table 4

**Supplementary Table 6.** Division times are longer in cells with aberrant PPBs in the *dcd3a-2 dcd3b* double mutant. Data was obtained from 8 *dcd3a-2 dcd3b* plants. Wilcoxon rank sum test with bonferroni correction 0.05/2 = 0.025 was performed to compare uneven and one-sided PPBs to cells with normal PPBs. * N <0.025 NS >0.025

| PPB Type | Median Metaphase Time (mins ± SE) | Wilcoxon test with bonferroni adj. p-value | Median Telophase Time (mins ± SE) | Wilcoxon test with bonferroni adj. p-value | Median Total Time (mins ± SE) | Wilcoxon test with bonferroni adj. p-value |
| --- | --- | --- | --- | --- | --- | --- |
| Normal (n=17 cells) | 30 ± 8 | | 40 ± 4 | | 90 ± 9 | |
| Uneven (n=23 cells) | 30 ± 6 | 0.26<br>NS | 60 ± 7 | 0.017<br>* | 110 ± 9 | 0.08<br>NS |
| One-Sided (n=7 cells) | 70 ± 20 | 0.06<br>NS | 70 ± 9 | 0.016<br>* | 160 ± 20 | 0.017<br>* |

**Supplementary Table 7.**
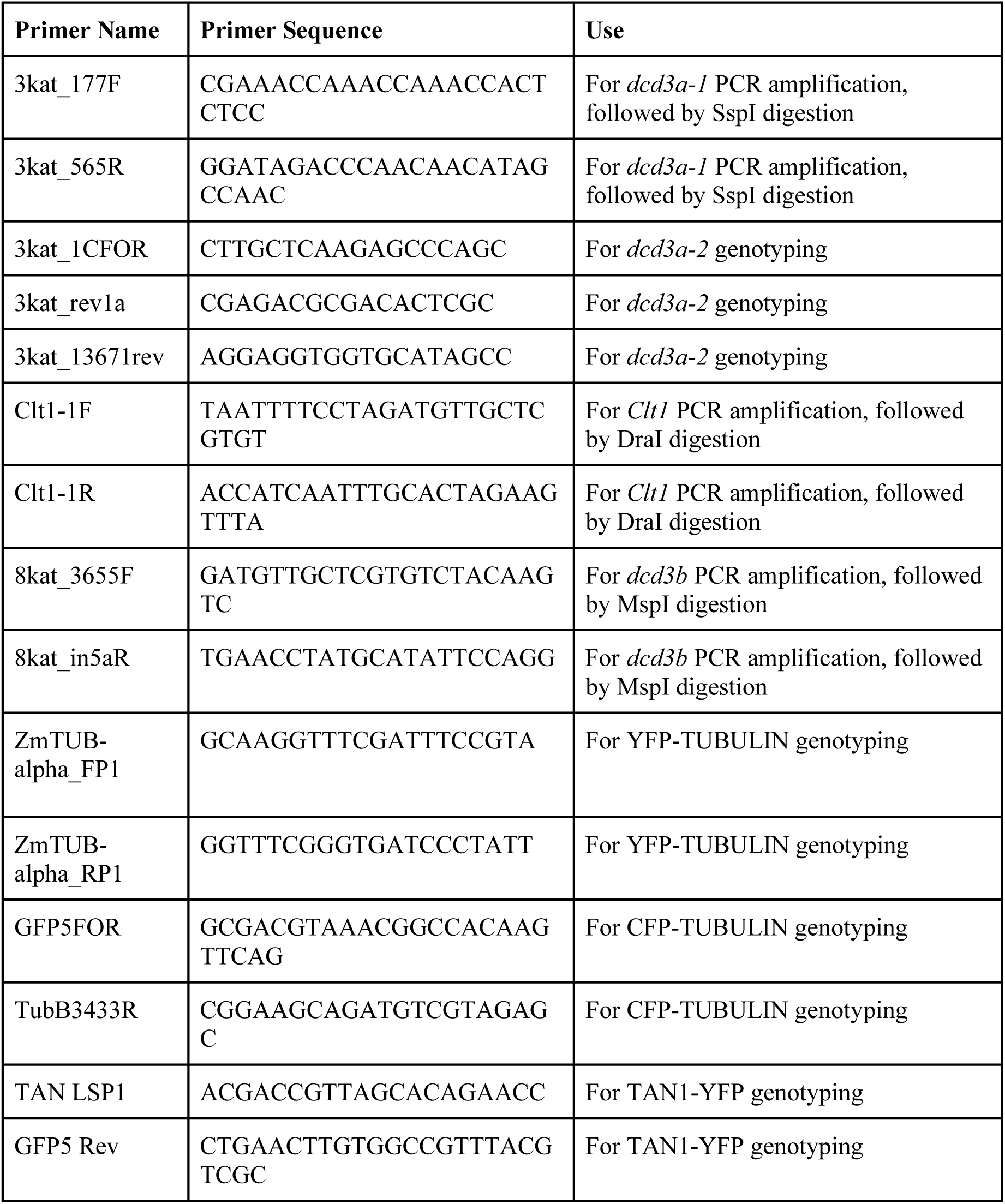
Primers used in this study.

## Notes

### Competing Interest Statement

The authors have declared no competing interest.

## References

1. Stoppin-Mellet, V. et al. Arabidopsis katanin binds microtubules using a multimeric microtubule-binding domain. Plant Physiol. Biochem. 45, 867–877 (2007).

2. McNally, F. J. & Vale, R. D. Identification of katanin, an ATPase that severs and disassembles stable microtubules. Cell 75, 419–429 (1993).

3. Hartman, J. J. & Vale, R. D. Microtubule disassembly by ATP-dependent oligomerization of the AAA enzyme katanin. Science 286, 782–785 (1999).

4. Hartman, J. J. et al. Katanin, a microtubule-severing protein, is a novel AAA ATPase that targets to the centrosome using a WD40-containing subunit. Cell 93, 277–287 (1998).

5. Wang, C. et al. KTN80 confers precision to microtubule severing by specific targeting of katanin complexes in plant cells. EMBO J. 36, 3435–3447 (2017).

6. Zhang, Q., Fishel, E., Bertroche, T. & Dixit, R. Microtubule severing at crossover sites by katanin generates ordered cortical microtubule arrays in Arabidopsis. Curr. Biol. 23, 2191–2195 (2013).

7. Lindeboom, J. J. et al. A mechanism for reorientation of cortical microtubule arrays driven by microtubule severing. Science 342, 1245533 (2013).

8. Deinum, E. E., Tindemans, S. H., Lindeboom, J. J. & Mulder, B. M. How selective severing by katanin promotes order in the plant cortical microtubule array. Proceedings of the National Academy of Sciences 114, 6942–6947 (2017).

9. Zehr, E. A., Szyk, A., Szczesna, E. & Roll-Mecak, A. Katanin Grips the β-Tubulin Tail through an Electropositive Double Spiral to Sever Microtubules. Dev. Cell 52, 118–131.e6 (2020).

10. Lu, C., Srayko, M. & Mains, P. E. The Caenorhabditis elegans microtubule-severing complex MEI-1/MEI-2 katanin interacts differently with two superficially redundant beta-tubulin isotypes. Mol. Biol. Cell 15, 142–150 (2004).

11. Bichet, A., Desnos, T., Turner, S., Grandjean, O. & Höfte, H. BOTERO1 is required for normal orientation of cortical microtubules and anisotropic cell expansion in Arabidopsis. Plant J. 25, 137– 148 (2001).

12. Bouquin, T., Mattsson, O., Naested, H., Foster, R. & Mundy, J. The Arabidopsis lue1 mutant defines a katanin p60 ortholog involved in hormonal control of microtubule orientation during cell growth. J. Cell Sci. 116, 791–801 (2003).

13. Komorisono, M. et al. Analysis of the rice mutant dwarf and gladius leaf 1. Aberrant katanin-mediated microtubule organization causes up-regulation of gibberellin biosynthetic genes independently of gibberellin signaling. Plant Physiol. 138, 1982–1993 (2005).

14. Webb, M., Jouannic, S., Foreman, J., Linstead, P. & Dolan, L. Cell specification in the Arabidopsis root epidermis requires the activity of ECTOPIC ROOT HAIR 3 – a katanin-p60 protein. Development 129, 123–131 (2002).

15. Wang, H. et al. CsKTN1 for a katanin p60 subunit is associated with the regulation of fruit elongation in cucumber (Cucumis sativus L.). Theor. Appl. Genet. (2021) doi:10.1007/s00122-021-03833-y.

16. McClinton, R. S., Chandler, J. S. & Callis, J. cDNA isolation, characterization, and protein intracellular localization of a katanin-like p60 subunit from Arabidopsis thaliana. Protoplasma 216, 181–190 (2001).

17. Burk, D. H., Liu, B., Zhong, R., Morrison, W. H. & Ye, Z. H. A katanin-like protein regulates normal cell wall biosynthesis and cell elongation. Plant Cell 13, 807–827 (2001).

18. Burk, D. H. & Ye, Z.-H. Alteration of oriented deposition of cellulose microfibrils by mutation of a katanin-like microtubule-severing protein. Plant Cell 14, 2145–2160 (2002).

19. Zhong, R., Burk, D. H., Morrison, W. H., 3rd & Ye, Z.-H. A kinesin-like protein is essential for oriented deposition of cellulose microfibrils and cell wall strength. Plant Cell 14, 3101–3117 (2002).

20. Komis, G. et al. Katanin Effects on Dynamics of Cortical Microtubules and Mitotic Arrays in Arabidopsis thaliana Revealed by Advanced Live-Cell Imaging. Front. Plant Sci. 8, 866 (2017).

21. Luptovčiak, I., Komis, G., Takáč, T., Ovečka, M. & Šamaj, J. Katanin: A Sword Cutting Microtubules for Cellular, Developmental, and Physiological Purposes. Front. Plant Sci. 8, 1982 (2017).

22. Yagi, N. et al. An anchoring complex recruits katanin for microtubule severing at the plant cortical nucleation sites. Nat. Commun. 12, 3687 (2021).

23. Belteton, S. A. et al. Real-time conversion of tissue-scale mechanical forces into an interdigitated growth pattern. Nat Plants 7, 826–841 (2021).

24. Peaucelle, A., Wightman, R. & Höfte, H. The Control of Growth Symmetry Breaking in the Arabidopsis Hypocotyl. Curr. Biol. 25, 1746–1752 (2015).

25. Uyttewaal, M. et al. Mechanical stress acts via katanin to amplify differences in growth rate between adjacent cells in Arabidopsis. Cell 149, 439–451 (2012).

26. Pickett-Heaps, J. D. & Northcote, D. H. Organization of microtubules and endoplasmic reticulum during mitosis and cytokinesis in wheat meristems. J. Cell Sci. 1, 109–120 (1966).

27. Smertenko, A. et al. Plant Cytokinesis: Terminology for Structures and Processes. Trends Cell Biol. 27, 885–894 (2017).

28. Panteris, E., Adamakis, I.-D. S., Voulgari, G. & Papadopoulou, G. A role for katanin in plant cell division: microtubule organization in dividing root cells of fra2 and lue1Arabidopsis thaliana mutants. Cytoskeleton 68, 401–413 (2011).

29. Attrill, S. T. & Dolan, L. KATANIN-mediated microtubule severing is required for MTOC organisation and function in Marchantia polymorpha. Development 151, (2024).

30. Panteris, E. & Adamakis, I.-D. S. Aberrant microtubule organization in dividing root cells of p60-katanin mutants. Plant Signal. Behav. 7, 16–18 (2012).

31. Sasaki, T. et al. A Novel Katanin-Tethering Machinery Accelerates Cytokinesis. Curr. Biol. 29, 4060–4070.e3 (2019).

32. Schmidt, S. & Smertenko, A. Identification and characterization of the land-plant-specific microtubule nucleation factor MACET4. J. Cell Sci. 132, (2019).

33. Ovečka, M. et al. Spatiotemporal Pattern of Ectopic Cell Divisions Contribute to Mis-Shaped Phenotype of Primary and Lateral Roots of katanin1 Mutant. Front. Plant Sci. 11, 734 (2020).

34. Panteris, E., Kouskouveli, A., Pappas, D. & Adamakis, I.-D. S. Cytokinesis in fra2 Arabidopsis thaliana p60-Katanin Mutant: Defects in Cell Plate/Daughter Wall Formation. Int. J. Mol. Sci. (2021) doi:10.20944/preprints202101.0226.v1.

35. Hazelwood, O. S., Orr, J. M. & Ashraf, M. A. Nuclear positioning and cell division site specification in plants. J. Exp. Bot. eraf241 (2025) doi:10.1093/jxb/eraf241.

36. Moukhtar, J. et al. Cell geometry determines symmetric and asymmetric division plane selection in Arabidopsis early embryos. PLoS Comput. Biol. 15, e1006771 (2019).

37. Gumber, H. K. et al. MLKS2 is an ARM domain and F-actin-associated KASH protein that functions in stomatal complex development and meiotic chromosome segregation. Nucleus 10, 144–166 (2019).

38. Ashraf, M. A., Liu, L. & Facette, M. R. A polarized nuclear position specifies the correct division plane during maize stomatal development. Plant Physiol. (2023) doi:10.1093/plphys/kiad329.

39. Wada, M. Nuclear movement and positioning in plant cells. Semin. Cell Dev. Biol. (2017) doi:10.1016/j.semcdb.2017.10.001.

40. Facette, M. R., Rasmussen, C. G. & Van Norman, J. M. A plane choice: coordinating timing and orientation of cell division during plant development. Curr. Opin. Plant Biol. 47, 47–55 (2019).

41. Kimata, Y. et al. Cytoskeleton dynamics control the first asymmetric cell division in Arabidopsis zygote. Proceedings of the National Academy of Sciences 113, 14157–14162 (2016).

42. Muroyama, A., Gong, Y. & Bergmann, D. C. Opposing, Polarity-Driven Nuclear Migrations Underpin Asymmetric Divisions to Pattern Arabidopsis Stomata. Curr. Biol. (2020) doi:10.1016/j.cub.2020.08.100.

43. Frank, M. J., Cartwright, H. N. & Smith, L. G. Three Brick genes have distinct functions in a common pathway promoting polarized cell division and cell morphogenesis in the maize leaf epidermis. Development 130, 753–762 (2003).

44. Panteris, E., Apostolakos, P. & Galatis, B. Cytoskeletal asymmetry in Zea mays subsidiary cell mother cells: a monopolar prophase microtubule half-spindle anchors the nucleus to its polar position. Cell Motil. Cytoskeleton 63, 696–709 (2006).

45. Murata, T. & Wada, M. Effects of centrifugation on preprophase-band formation in Adiantum protonemata. Planta 183, 391–398 (1991).

46. Wright, A. J., Gallagher, K. & Smith, L. G. discordia1 and alternative discordia1 function redundantly at the cortical division site to promote preprophase band formation and orient division planes in maize. Plant Cell 21, 234–247 (2009).

47. Uyehara, A. N. et al. De novo TANGLED1 recruitment from the phragmoplast to aberrant cell plate fusion sites in maize. J. Cell Sci. 137, (2024).

48. Gallagher, K. & Smith, L. G. discordia mutations specifically misorient asymmetric cell divisions during development of the maize leaf epidermis. Development 126, 4623–4633 (1999).

49. Nan, Q. et al. The OPAQUE1/DISCORDIA2 myosin XI is required for phragmoplast guidance during asymmetric cell division in maize. Plant Cell 35, 2678–2693 (2023).

50. Wang, G. et al. Opaque1 encodes a myosin XI motor protein that is required for endoplasmic reticulum motility and protein body formation in maize endosperm. Plant Cell 24, 3447–3462 (2012).

51. Zebosi, B. et al. Recessive antimorph alleles reveal novel functions of the OPAQUE1 myosin XI in maize. bioRxiv 2025.06. 26.661838 (2025) doi:10.1101/2025.06.26.661838.

52. M. G. Neuffer and K. A. Sheridan. Dominant mutants from EMS-treated pollen. Maize Genetics Cooperation News Letter 59–60 (1977).

53. Harper, L., Gardiner, J., Andorf, C. & Lawrence, C. J. MaizeGDB: The maize genetics and genomics database. Methods Mol. Biol. 1374, 187–202 (2016).

54. Andorf, C. M. et al. MaizeGDB update: new tools, data and interface for the maize model organism database. Nucleic Acids Res. 44, D1195–201 (2016).

55. Riglet, L. et al. KATANIN-dependent mechanical properties of the stigmatic cell wall mediate the pollen tube path in Arabidopsis. Elife 9, (2020).

56. Hamada, T. Microtubule organization and microtubule-associated proteins in plant cells. Int. Rev. Cell Mol. Biol. 312, 1–52 (2014).

57. Bouquin, T., Mattsson, O., Naested, H., Foster, R. & Mundy, J. The Arabidopsis lue1 mutant defines a katanin p60 ortholog involved in hormonal control of microtubule orientation during cell growth. J. Cell Sci. 116, 791–801 (2003).

58. Bichet, A., Desnos, T., Turner, S., Grandjean, O. & Höfte, H. BOTERO1 is required for normal orientation of cortical microtubules and anisotropic cell expansion in Arabidopsis: BOTERO1 is required for microtubule alignment. Plant J. 25, 137–148 (2008).

59. Komorisono, M. et al. Analysis of the rice mutant dwarf and gladius leaf 1. Aberrant katanin-mediated microtubule organization causes up-regulation of gibberellin biosynthetic genes independently of gibberellin signaling. Plant Physiol. 138, 1982–1993 (2005).

60. Sun, X. et al. Altered expression of maize PLASTOCHRON1 enhances biomass and seed yield by extending cell division duration. Nat. Commun. 8, 14752 (2017).

61. Sprangers, K., Avramova, V. & Beemster, G. T. S. Kinematic analysis of cell division and expansion: Quantifying the cellular basis of growth and sampling developmental zones in Zea mays leaves. J. Vis. Exp. (2016) doi:10.3791/54887.

62. Nelissen, H. et al. A local maximum in gibberellin levels regulates maize leaf growth by spatial control of cell division. Curr. Biol. 22, 1183–1187 (2012).

63. R Jones, A., et al. Cell-size dependent progression of the cell cycle creates homeostasis and flexibility of plant cell size. Nat. Commun. 8, 15060 (2017).

64. D’Ario, M. et al. Cell size controlled in plants using DNA content as an internal scale. Science 372, 1176–1181 (2021).

65. Willis, L. et al. Cell size and growth regulation in the Arabidopsis thaliana apical stem cell niche. Proc. Natl. Acad. Sci. U. S. A. 113, E8238–E8246 (2016).

66. Panteris, E., Diannelidis, B.-E. & Adamakis, I.-D. S. Cortical microtubule orientation in Arabidopsis thaliana root meristematic zone depends on cell division and requires severing by katanin. *J*. Biol. Res. 25, 12 (2018).

67. Desvoyes, B., Arana-Echarri, A., Barea, M. D. & Gutierrez, C. A comprehensive fluorescent sensor for spatiotemporal cell cycle analysis in Arabidopsis. Nature Plants 6, 1330–1334 (2020).

68. Fan, Y., Burkart, G. M. & Dixit, R. The Arabidopsis SPIRAL2 Protein Targets and Stabilizes Microtubule Minus Ends. Curr. Biol. 28, 987–994.e3 (2018).

69. Li, Y. et al. ABNORMAL SHOOT 6 interacts with KATANIN 1 and SHADE AVOIDANCE 4 to promote cortical microtubule severing and ordering in Arabidopsis. J. Integr. Plant Biol. 63, 646– 661 (2021).

70. Wang, J. et al. Brassinosteroid signals cooperate with katanin-mediated microtubule severing to control stamen filament elongation. EMBO J. e111883 (2022) doi:10.15252/embj.2022111883.

71. Song, G. et al. The Rice SPOTTED LEAF4 (SPL4) Encodes a Plant Spastin That Inhibits ROS Accumulation in Leaf Development and Functions in Leaf Senescence. Front. Plant Sci. 9, 1925 (2018).

72. Zhang, T., Zhao, S.-H., Wang, Y. & He, Y. FIGL1 coordinates with dosage-sensitive BRCA2 in modulating meiotic recombination in maize. J. Integr. Plant Biol. 65, 2107–2121 (2023).

73. Schaefer, E. et al. The preprophase band of microtubules controls the robustness of division orientation in plants. Science 356, 186–189 (2017).

74. Zhang, Y., Iakovidis, M. & Costa, S. Control of patterns of symmetric cell division in the epidermal and cortical tissues of the Arabidopsis root. Development 143, 978–982 (2016).

75. Pietra, S., Lang, P. & Grebe, M. SABRE is required for stabilization of root hair patterning in Arabidopsis thaliana. Physiol. Plant. 153, 440–453 (2015).

76. Pietra, S. et al. Arabidopsis SABRE and CLASP interact to stabilize cell division plane orientation and planar polarity. Nat. Commun. 4, 2779 (2013).

77. Rony, R. M. I. K., Campos, R., Pérez-Henríquez, P. & Van Norman, J. M. Outward askew endodermal cell divisions reveal INFLORESCENCE AND ROOT APICES RECEPTOR KINASE functions in division orientation. Plant Physiol. 196, 2251–2262 (2024).

78. Flanders, D. J., Rawlins, D. J., Shaw, P. J. & Lloyd, C. W. Nucleus-associated microtubules help determine the division plane of plant epidermal cells: avoidance of four-way junctions and the role of cell geometry. J. Cell Biol. 110, 1111–1122 (1990).

79. Boruc, J., Zhou, X. & Meier, I. Dynamics of the Plant Nuclear Envelope and Nuclear Pore. Plant Physiol. 158, 78–86 (2012).

80. Wick, S. M. & Duniec, J. Immunofluorescence microscopy of tubulin and microtubule arrays in plant cells. I. Preprophase band development and concomitant appearance of nuclear envelope-associated tubulin. J. Cell Biol. 97, 235–243 (1983).

81. Marcus, A. I., Li, W., Ma, H. & Cyr, R. J. A kinesin mutant with an atypical bipolar spindle undergoes normal mitosis. Mol. Biol. Cell 14, 1717–1726 (2003).

82. Ambrose, J. C. & Cyr, R. Mitotic spindle organization by the preprophase band. Mol. Plant 1, 950– 960 (2008).

83. Ludwig, Y., Zhang, Y. & Hochholdinger, F. The maize (Zea mays L.) AUXIN/INDOLE-3-ACETIC ACID gene family: phylogeny, synteny, and unique root-type and tissue-specific expression patterns during development. PLoS One 8, e78859 (2013).

84. Draves, M. A., Muench, R. L., Lang, M. G. & Kelley, D. R. Maize seedling growth and hormone response assays using the rolled towel method. Curr. Protoc. 2, e562 (2022).

85. Candela, H., Johnston, R., Gerhold, A., Foster, T. & Hake, S. The milkweed pod1 gene encodes a KANADI protein that is required for abaxial/adaxial patterning in maize leaves. Plant Cell 20, 2073– 2087 (2008).

86. Mills, A., Allsman, L., Leon, S. & Rasmussen, C. Using seed chipping to genotype maize kernels. Bio Protoc. 10, (2020).

87. Rasmussen, C. G. Using Live-Cell Markers in Maize to Analyze Cell Division Orientation and Timing. Methods Mol. Biol. 1370, 209–225 (2016).

88. Thévenaz, P. StackReg: an ImageJ plugin for the recursive alignment of a stack of images. *Biomedical Imaging Group*, Swiss Federal Institute of Technology Lausanne 2012, (1998).

89. Domander, R., Felder, A. A. & Doube, M. BoneJ2 -refactoring established research software. Wellcome Open Res. 6, 37 (2021).

90. Boudaoud, A. et al. FibrilTool, an ImageJ plug-in to quantify fibrillar structures in raw microscopy images. Nat. Protoc. 9, 457–463 (2014).

91. Martinez, J. C. & Wang, K. A sterilization protocol for field-harvested maize mature seed used for in vitro culture and genetic transformation. Maize Genetics Cooperation Newsletter 83, (2009).

92. Farrow, J., Bellinger, M. A. & Rasmussen, C. G. In vitro Conditions for Dark Growth and Analysis of Maize Seedlings. Bio-protocol 10, e3555–e3555 (2020).

93. Nelissen, H. et al. Kinematic analysis of cell division in leaves of mono- and dicotyledonous species: a basis for understanding growth and developing refined molecular sampling strategies. Methods Mol. Biol. 959, 247–264 (2013).

94. Allsman, L. A., Dieffenbacher, R. N. & Rasmussen, C. G. Glue impressions of maize leaves and their use in classifying mutants. Bio-protocol 9, e3209 (2019).

95. Wickham, H., Vaughan, D. & Girlich, M. tidyr: Tidy Messy Data. tidyr: Tidy Messy Data. R package version 1.3.1 https://tidyr.tidyverse.org (2024).

96. Wickham, H. ggplot2: Elegant Graphics for Data Analysis. (Springer-Verlag New York, 2016).

97. Dawson, C. ggprism: A ‘ggplot2’ Extension Inspired by ‘GraphPad Prism’. https://cran.r-project.org/package=ggprism; doi: 10.5281/zenodo.4556067 (2024).

98. Hvitfeldt, E. paletteer: Comprehensive Collection of Color Palettes. https://github.com/EmilHvitfeldt/paletteer (2021).

99. Wickham, H. stringr: Simple, Consistent Wrappers for Common String Operations. R package version 1.5.1. https://github.com/tidyverse/stringr; https://stringr.tidyverse.org (2023).

